# Genomic insights into rapid diversification and adaptive introgression in the earthworm *Amynthas aspergillum*

**DOI:** 10.64898/2026.02.13.704776

**Authors:** Yangmei Qin, Zeng Peng, Yongsheng Mo, Deqiang Shi, Chunhua He, Jun Liu, Weiguan Zhou, Yuantao Zheng, Xiuzhi Li, Bo Lu, Mei Lin, Kaitai Yang, Lixin Peng

## Abstract

How species diversify in highly fragmented landscapes while maintaining genetic connectivity remains a central question in evolutionary biology. The megascolecid earthworm *Amynthas aspergillum* exemplifies this paradox, showing deeply divergent mitochondrial lineages despite extensive nuclear gene flow across southern China. We conducted whole-genome resequencing across its core distribution in the Guangxi karst region to elucidate the population structure and processes underlying this cytonuclear discordance. Genome-wide analyses identified three genetically distinct groups shaped by fine-scale karst isolation, alongside pervasive historical and contemporary introgression. Demographic modeling revealed rapid diversification during the Late Pleistocene (∼0.14LJMa) with pronounced fluctuations in effective population size. Although isolation increased inbreeding and mutational load in some lineages, introgression persisted even among deeply divergent mitochondrial clades. Notably, several introgressed regions showed strong signatures of positive selection, including genes involved in mitochondrial function and metabolism.Together, these results demonstrate that gene flow can simultaneously mitigate the genomic costs of isolation and promote functional innovation via adaptive introgression, providing a genomic framework for diversification with gene flow in subterranean invertebrates inhabiting fragmented karst landscapes.

## Introduction

Earthworms are foundational ecosystem engineers that regulate terrestrial ecosystem functioning through their effects on soil structure, nutrient cycling, and hydrological processes^1^. Among megascolecid earthworms, *Amynthas aspergillum* (Perrier, 1872) is a dominant species widely distributed across the subtropical landscapes of Southern China, particularly within the karst regions of Guangxi^2^. In these environments, *A. aspergillum* plays a critical role in soil water regulation and can reduce surface evaporation by more than 60% in lateritic red soils^3^. Beyond its ecological functions, *A. aspergillum* is also a premier medicinal resource and one of the few earthworm species formally recognized in the Chinese Pharmacopoeia for its diverse pharmacological properties^4,5^

A defining feature of *Amynthas aspergillum (A. aspergillum)* is its complex evolutionary history shaped by the fragmented topography of the South China karst. Phylogeographic analyses based on multilocus mitochondrial markers (*COI, COII*, and 16S rRNA) have revealed exceptionally high genetic diversity, comprising six deeply divergent mitochondrial clades (Clades I–VI) that originated in the Guangxi Basin during the Pliocene (∼4.61 Ma) ^6^. Paradoxically, despite these ancient maternal splits, mitochondrial clades of *A. aspergillum* show extensive geographic overlap and only weak phylogeographic structure^6^, suggesting substantial nuclear connectivity among lineages and resulting in pronounced cytonuclear discordance. While cytonuclear discordance has been documented across a wide array of taxa^7^, its evolutionary significance in subterranean systems remains poorly understood, contrasting with a growing body of phylogenomic and population genomic evidence for widespread genomic discordance across diverse evolutionary lineages^8,9^. This empirical gap raises a fundamental evolutionary question: how can deeply divergent mitochondrial lineages persist over extended evolutionary timescales in highly fragmented subterranean landscapes, despite extensive nuclear introgression that maintains genomic connectivity?

Recent genomic advances, including the assembly of a telomere-to-telomere (T2T) reference genome^10^, have provided a complete chromosomal framework for *A. aspergillum*. While this resource offers unprecedented resolution of genome architecture, it provides limited insight into population-level evolutionary dynamics, including how genetic diversity is structured, maintained, and reshaped across heterogeneous landscapes. Resolving these processes is essential for understanding the mechanisms underlying the ecological resilience and long-term persistence of this species.

In this study, we provide a high-resolution genomic framework to resolve this paradox by performing whole-genome resequencing of *A. aspergillum* across its primary distribution range in Guangxi. By integrating the T2T reference genome with demographic inference and introgression analyses, we explicitly test a model of speciation with gene flow in a highly fragmented karst landscape. Specifically, we aim to (i) reconstruct the tempo and mode of rapid diversification within the Guangxi karst system; (ii) quantify the extent and directionality of nuclear introgression among divergent mitochondrial clades; and (iii) evaluate the role of adaptive introgression in shaping functional genomic variation.

Our results reveal that, rather than acting solely as a barrier to divergence, gene flow —particularly in the form of adaptive introgression—has contributed to both genomic cohesion and functional innovation, while partially mitigating the accumulation of deleterious variation in isolated lineages. These findings establish *A. aspergillum* as a powerful model for understanding how fragmented landscapes simultaneously promote lineage persistence and evolutionary connectivity, offering broader insights into invertebrate evolution under complex topographic constraints.

## Results

### Fine-scale population structure and genetic diversity of A. aspergillum

To investigate the genomic landscape and population structure of *Amynthas aspergillum*, we performed high-depth whole-genome resequencing (WGS) of 48 individuals collected from 25 geographically representative sites across the Guangxi region, China (**Supplementary Fig. 1** & **Table S1**). The sequencing data demonstrated high mapping quality, with a median mapping rate of 98.56% and a median of 83.21% of the genome covered at depths greater than 5X (**Supplementary Fig. 2 & Table S2**). After stringent quality control (**Methods**) and alignment to a high-quality telomere-to-telomere (T2T) reference genome (746.95 Mb, 43 chromosomes)^10^, we identified approximately 10.7 million high-quality single nucleotide polymorphisms (SNPs) (**Table S3**). SNP heterozygosity ratios (the proportion of heterozygous sites among identified variants) were comparable between Group 1 (22.88% – 25.51%, mean = 24.32%) and Group 3 (23.46% – 25.23%, mean=24.19%), whereas Group 2 exhibited moderately lower values (18.04% – 22.67%, mean=21.55%) (**Table S4 & Table S5**). When normalized to the estimated genome size (746.95 Mb) ^10^, the genome-wide heterozygosity rates of these three core groups remained relatively low and stable, ranging from 0.18% to 0.29%. In contrast, individuals in the OutGroup showed higher SNP heterozygosity (mean = 30.21%) and an elevated genome-wide heterozygosity rate (up to 0.39%), consistent with their greater genetic divergence from the focal populations (**Table S4**).

Preliminary clustering based on mitochondrial genome analyses identified three individuals (B28, L14, and Y13) with close phylogenetic affinity to *A. robustus* (**Supplementary Fig. 3**). Although these samples exhibited relatively high mapping rates (95.26%, 84.90%, and 98.14%), their genome coverage at ≥1× depth was lower (67.91%, 53.03%, and 81.78%, respectively) than that of the core individuals (**Table S2**). These individuals were therefore designated as an outgroup for all subsequent nuclear genomic analyses (**Fig. 1A**). Although the lowest cross-validation (CV) error in ADMIXTURE^11^ was observed at K = 2 (**Supplementary Fig. 4**), we selected K = 4 for downstream analyses because it better matched the genetic structure revealed by principal component analysis (PCA) (**Fig. 1B**). At K = 4, lineage divergence was resolved with greater clarity, consistent with patterns along the major principal components, whereas lower K values failed to capture this finer-scale structure (**Supplementary Fig. 5**).

**Figure 1.**
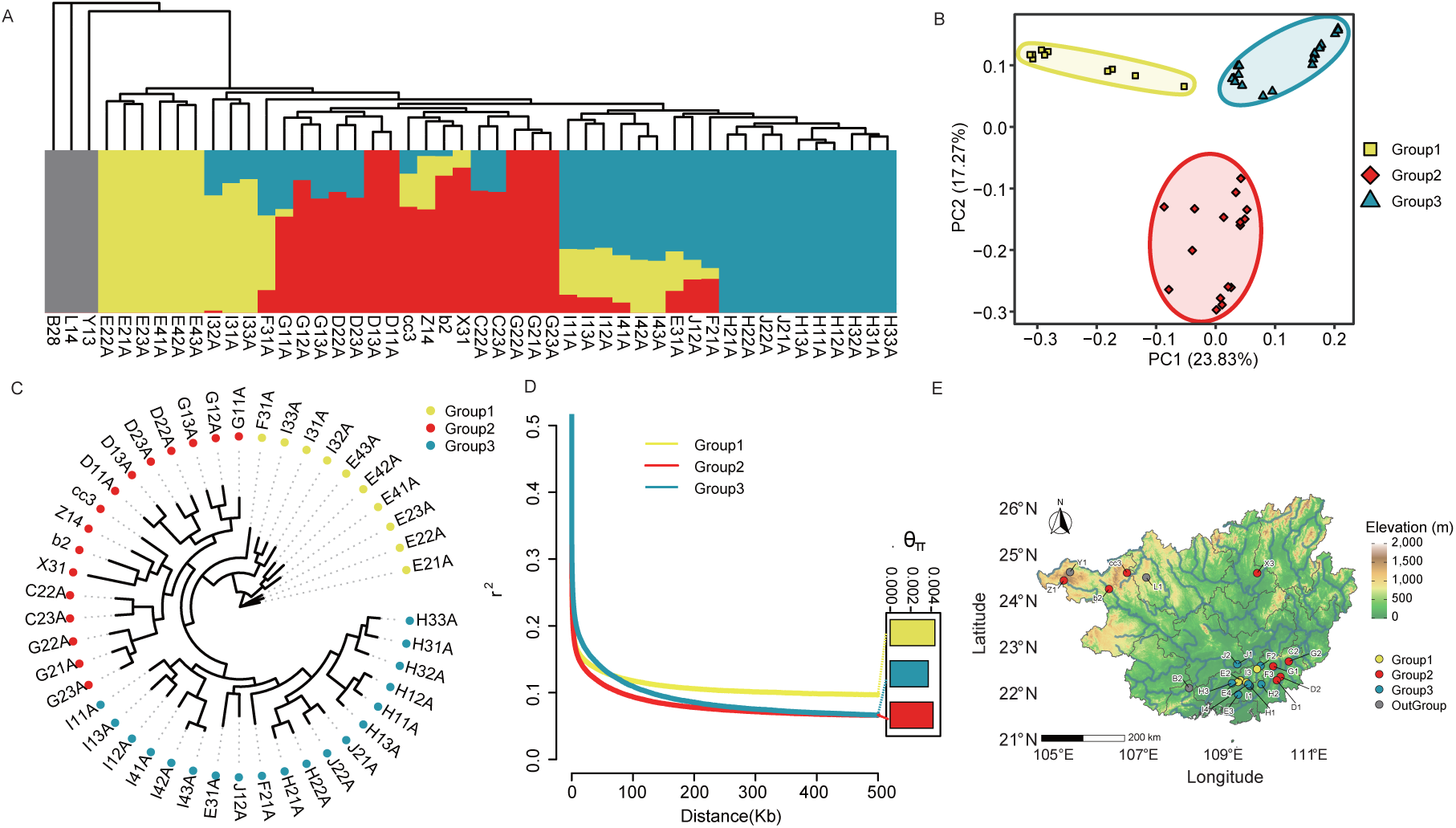
Geographic distribution and genomic landscape of earthworm populations. (A) Genetic structure and admixture analysis. Maximum likelihood phylogenetic tree (top) and ADMIXTURE clustering (bottom) based on genome-wide SNPs. Samples are classified into three primary genetic clusters: Group 1 (yellow), Group 2 (red), and Group 3 (blue), with *B28*, *L14*, and *Y13* serving as the outgroup (grey). (B) Principal component analysis (PCA). Projection of individuals onto the first two principal components. PC1 and PC2 explain 23.83% and 17.27% of the total genetic variance, respectively. Symbols represent the three groups, consistent with the colors in a. (C) Nuclear phylogenomic reconstruction. A circularized neighbor-joining tree illustrating the relationships among sampled individuals. Tips are color-coded by their respective genetic groups. (D) Patterns of linkage disequilibrium (LD) and nucleotide diversity. Decay of LD (measured as r^2) against physical distance (kb) for the three populations. The inset bar plot displays the nucleotide diversity (pi) for each group, indicating varied levels of genetic variation across populations. (E) Topographic distribution of sampling sites. Sampling locations are overlaid on a Digital Elevation Model (DEM). Point colors denote the genetic groups identified in b. Group 2 (red) is situated adjacent to the Group 1–3 complex; notably, a subset of Group 2 individuals (including samples cc3, Z1, b2, and X3) occurs in high-elevation or high-latitude regions characterized by rugged topography.

ADMIXTURE analysis at K = 4 revealed largely stable ancestral components within each group, with limited admixture in several individuals, indicative of historical or ongoing gene flow among divergent lineages (**Fig. 1A**). Population structure analyses consistently identified three distinct genetic clusters among the 45 *A. aspergillum individuals*. PCA clearly separated individuals into three groups—Group 1 (yellow), Group 2 (red), and Group 3 (blue)—with the first two principal components (PC1 and PC2) explaining 23.83% and 17.27% of the total genetic variance, respectively (**Fig. 1B**). These groupings were further corroborated by maximum likelihood (ML) and neighbor-joining (NJ) phylogenetic analyses (**Methods**), both of which recovered three well-supported monophyletic clades corresponding to the PCA-defined clusters (**Fig. 1C**). The final sample composition comprised 10 individuals in Group 1, 16 in Group 2, and 19 in Group 3, with the remaining three samples assigned to an outgroup for comparative analyses.

To further characterize genetic differentiation among the three groups, we examined nucleotide diversity (π) and linkage disequilibrium (LD) patterns. Nucleotide diversity varied among groups, with Group 1 exhibiting the highest π, followed by Group 2 and Group 3 (**Fig. 1D & Table S5**). LD decay analyses revealed contrasting patterns (**Fig. 1D**). Group 2 showed the most rapid decay of r², whereas Group 1 maintained elevated r² values over longer physical distances. Group 3 displayed strong short-range LD followed by a sharp decline with increasing distance. Notably, the elevated LD observed in Group 1 despite its relatively high nucleotide diversity suggests that factors beyond long-term effective population size—such as recent demographic fluctuations, population substructure, or selective processes—may have shaped its genomic architecture.

To further investigate the evolutionary forces shaping these populations, we calculated multiple neutrality and differentiation statistics (**Table S5**). All three groups exhibited positive Tajima’s D values (Group 1: 0.84; Group 2: 1.02; Group 3: 0.83), suggesting a relative deficit of rare alleles. In line with the individual-level analyses, Group 2 showed the lowest average heterozygosity (21.55%), whereas Group 1 and Group 3 maintained higher levels (24.32% and 24.19%, respectively), indicating comparatively reduced genetic diversity in Group 2. Pairwise differentiation among the three groups was moderate. The highest divergence was observed between Group 1 and Group 3 (FST = 0.1344), while Group 2 exhibited lower differentiation from both Group 1 (FST = 0.1045) and Group 3 (FST = 0.0835). In contrast, differentiation between each group and the OutGroup was substantially higher (FST = 0.3634–0.4324), reflecting pronounced genomic divergence between the core clusters and the reference population.

Despite the absence of obvious geographic barriers, the three genetic groups exhibited discernible spatial structuring across the study area (**Fig. 1E**). Group 1 (yellow) was mainly concentrated in the central region, whereas Group 3 (blue) was more frequently found toward peripheral areas. Group 2 (red) occurred in locations adjacent to both Groups 1 and 3 but showed limited spatial overlap with either, resulting in a largely juxtaposed spatial distribution.

Overall, these patterns indicate that distinct genetic boundaries can be maintained even without clear physical barriers. Such structuring may reflect the combined effects of landscape heterogeneity, historical demographic processes, and selective pressures, illustrating how complex subterranean environments can contribute to population differentiation across space.

### Pleistocene demographic dynamics and asymmetric introgression shape population structure in Amynthas aspergillum

To elucidate the temporal and evolutionary processes underlying population differentiation in *A. aspergillum*, we inferred a time-calibrated phylogeny using MCMCTree^12^ (**Fig. 2A; Methods**). Molecular clock analyses estimated the divergence between the Amynthas lineage and Eisenia at approximately 230 Ma, whereas the split between Eisenia and Capitella was dated to approximately 470 Ma (**Table S6**). These estimates are broadly consistent with previously reported fossil-calibrated divergence times for major annelid lineages^13,14^, supporting the reliability of our molecular dating framework. At a finer temporal scale, divergence time estimation indicated that *A. aspergillum* diverged from its sister species A. corticis at approximately 48 Ma, with a 95% highest posterior density (HPD) interval of 41–54 Ma, a timeframe corresponding to the middle Eocene (Paleogene). **(Fig. 2A; Table S6**).

**Figure 2.**
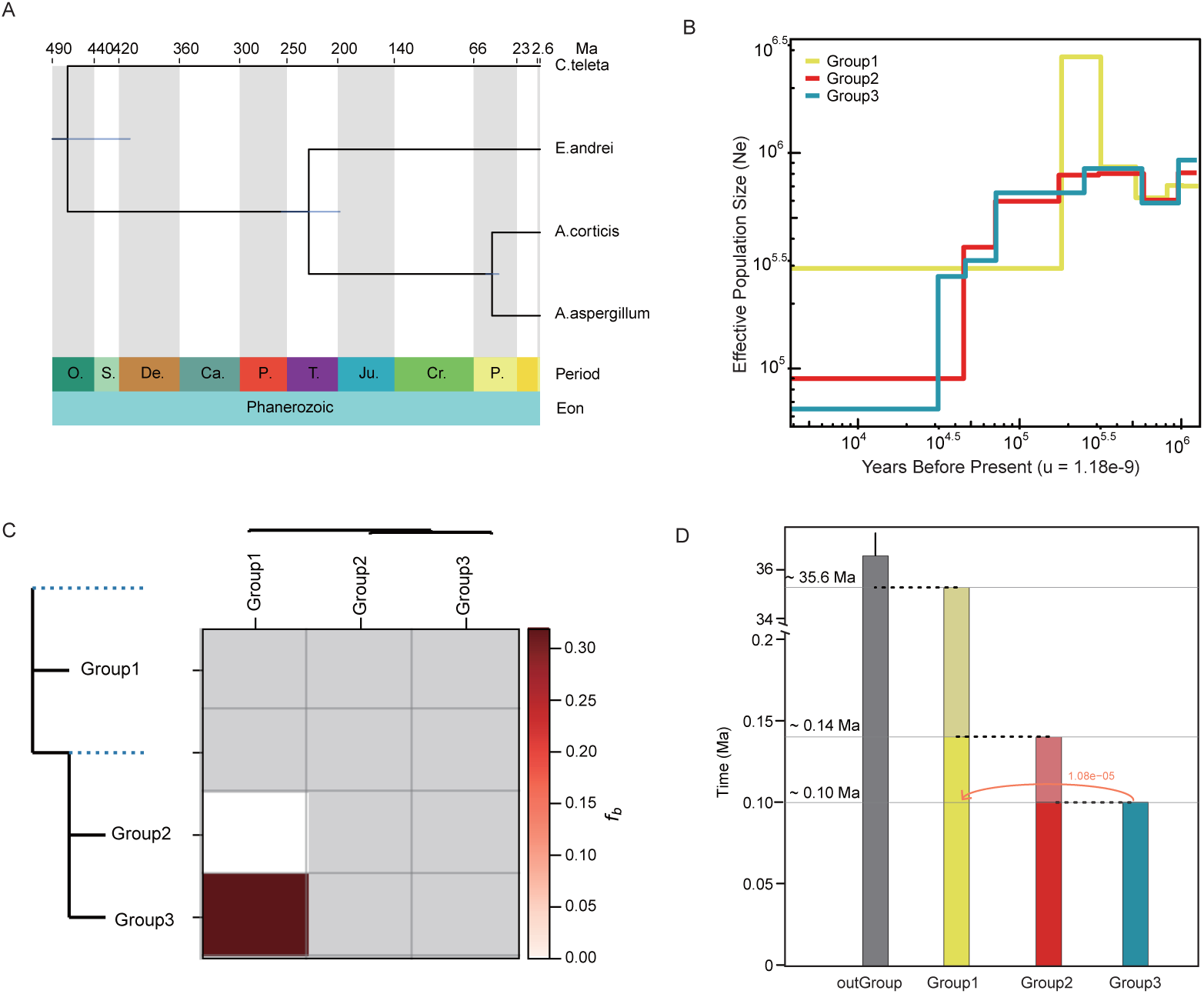
Demographic history and genome-wide introgression patterns. (A) Macroevolutionary divergence and time calibration. A time-calibrated phylogenetic tree showing the deep evolutionary relationships among species. Divergence times are indicated in millions of years (Ma) at the top, aligned with the geological periods and eons of the Phanerozoic. Blue bars at nodes indicate 95% highest posterior density (HPD) intervals for divergence estimates. (B) Dynamic changes in effective population size (Ne). Demographic history inferred by SMC++ across the three genetic groups (Group 1: yellow; Group 2: red; Group 3: blue). The y-axis represents the Ne on a log scale, and the x-axis represents the time in years before present (assuming a mutation rate u = 1.18*10^-9. All groups show a dramatic expansion approximately 10^5 to 10^5.5 years ago, followed by recent declines. (C) Patterns of f-branch (f_b_) statistics. Heatmap illustrating gene flow between specific lineages. The tree on the left represents the tested topology (Group1, Group2 and Group3). The color intensity of the squares corresponds to the f_b_ value, indicating the excess shared ancestry between lineages. Strong introgression signals are observed between the Group1 and Group3 lineages. (D) Coalescence-based divergence modeling and migration. A summary of demographic parameters inferred from fastsimcoal2. Bar heights represent the divergence times of lineages: the split between the outgroup and ingroups is anchored at ∼35.6 Ma, while internal lineage splits are estimated at ∼0.14 Ma and ∼0.10 Ma. The orange arrow indicates bidirectional gene flow from Group 3 to Group 1, with the migration rate (1.08*10^-5) labeled above.

Historical reconstructions of effective population size using SMC++^15^ reveal highly concordant demographic trajectories across the three genetic groups (**Fig. 2B & Methods**). All lineages maintained large effective population sizes during the Early to Middle Pleistocene, exhibiting a synchronous early Ne peak of approximately 1.20–1.33 × 10^6 between 1.39 and 1.64 Ma (**Table S7**). Following this shared expansion, Group 1 underwent a pronounced secondary increase in effective population size, reaching a peak of approximately 2.79 × 10^6 around 0.31 Ma, whereas Groups 2 and 3 showed a sustained decline in Ne thereafter. These demographic changes were followed by an overall trend of population contraction toward the present. Notably, recent effective population size estimates differ markedly among lineages: Group 1 retains a substantially higher near-present Ne (approximately 2.91 × 10^5 at 0.18 Ma) compared with Group 2 (8.99 × 10^4 at 0.04 Ma) and Group 3 (6.49 × 10^4 at 0.03 Ma). These differences are consistent with the elevated genome-wide nucleotide diversity and slower linkage disequilibrium decay detected in Group 1 relative to the other groups (**Fig. 1D; Table S7**).

Genome-wide f-branch (fb) statistics were further used to assess whether differential gene flow contributed to the contrasting demographic and diversity patterns observed among the three groups (**Methods**). A pronounced excess of shared ancestry (fb = 0.3185) indicating substantial historical introgression (**Fig. 2C & Table S8**). In contrast, Group 2 exhibits minimal evidence of genetic exchange with either lineage, suggesting a comparatively higher degree of reproductive or spatial isolation.

Demographic models implemented in fastsimcoal2^16^ integrate divergence timing and migration into a unified framework (**Fig. 2D**). Model-based inference indicates that the Guangxi ingroup diverged from the outgroup approximately 35.6 Ma, whereas diversification within the region occurred much more recently. Specifically, the split between Group 1 and the ancestor of Groups 2 and Group 3 was estimated at ∼0.14 Ma, followed by divergence between Groups 2 and 3 at ∼0.10 Ma. Consistent with the fb results, the best-supported model inferred significant asymmetric migration predominantly from Group 3 into Group 1 (migration rate = 1.08 × 10^−5), whereas migration in the opposite direction was negligible (estimated migration rate = 4.32 × 10^−8).

Together, these analyses reveal a pronounced temporal disparity in the evolutionary history of *A. aspergillum* in Guangxi. Although the lineage diverged from its closest relatives deep in time (∼35.6 Ma), genomic evidence indicates that diversification within the Guangxi populations occurred remarkably recently, during the Late Pleistocene (∼0.10–0.14 Ma). This shallow internal divergence coincides with highly heterogeneous demographic trajectories inferred from SMC++, with Group 1 experiencing a secondary population expansion while Groups 2 and 3 underwent sustained declines in effective population size. Patterns of historical gene flow further indicate that these demographic differences were accompanied by strongly lineage-specific connectivity, characterized by excess shared ancestry between Group 1 and Group 3 and predominantly unidirectional migration from Group 3 into Group 1. Collectively, these results suggest that recent population differentiation in *A. aspergillum* reflects the combined effects of rapid late-stage divergence, contrasting demographic histories, and uneven historical gene flow among lineages.

### Divergent genomic consequences under shared demographic histories

Despite broadly concordant Early to Middle Pleistocene demographic trajectories inferred across the three genetic groups (**Fig. 2B**), their genomes exhibit strikingly divergent signatures of inbreeding and mutational load. Comparative analyses based on runs of homozygosity (ROH) reveal pronounced disparities in inbreeding levels among lineages (**Fig. 3A & Methods).** The absence of long ROH (>5 Mb) across all groups suggests that elevated homozygosity is driven by historical population contraction rather than recent consanguineous mating **(Supplementary Fig. 6A**). Group 3 exhibits the highest ROH burden, with ROH-based inbreeding coefficients (F_ROH) reaching up to 0.1500 (e.g., individual H12A), whereas substantially lower values are observed in Group 1 (e.g., individual I31A, F_ROH = 0.0033) (**Supplementary Fig. 6B & Table S9**). This markedly elevated inbreeding in Group 3 is consistent with its reduced nucleotide diversity and slower decay of linkage disequilibrium (**Fig. 1D**). Although Group 2 also exhibits higher F_ROH values than Group 1, the extreme inbreeding levels observed in Group 3 point to a more severe history of restricted connectivity or persistently reduced effective population size in recent evolutionary history.

**Figure 3.**
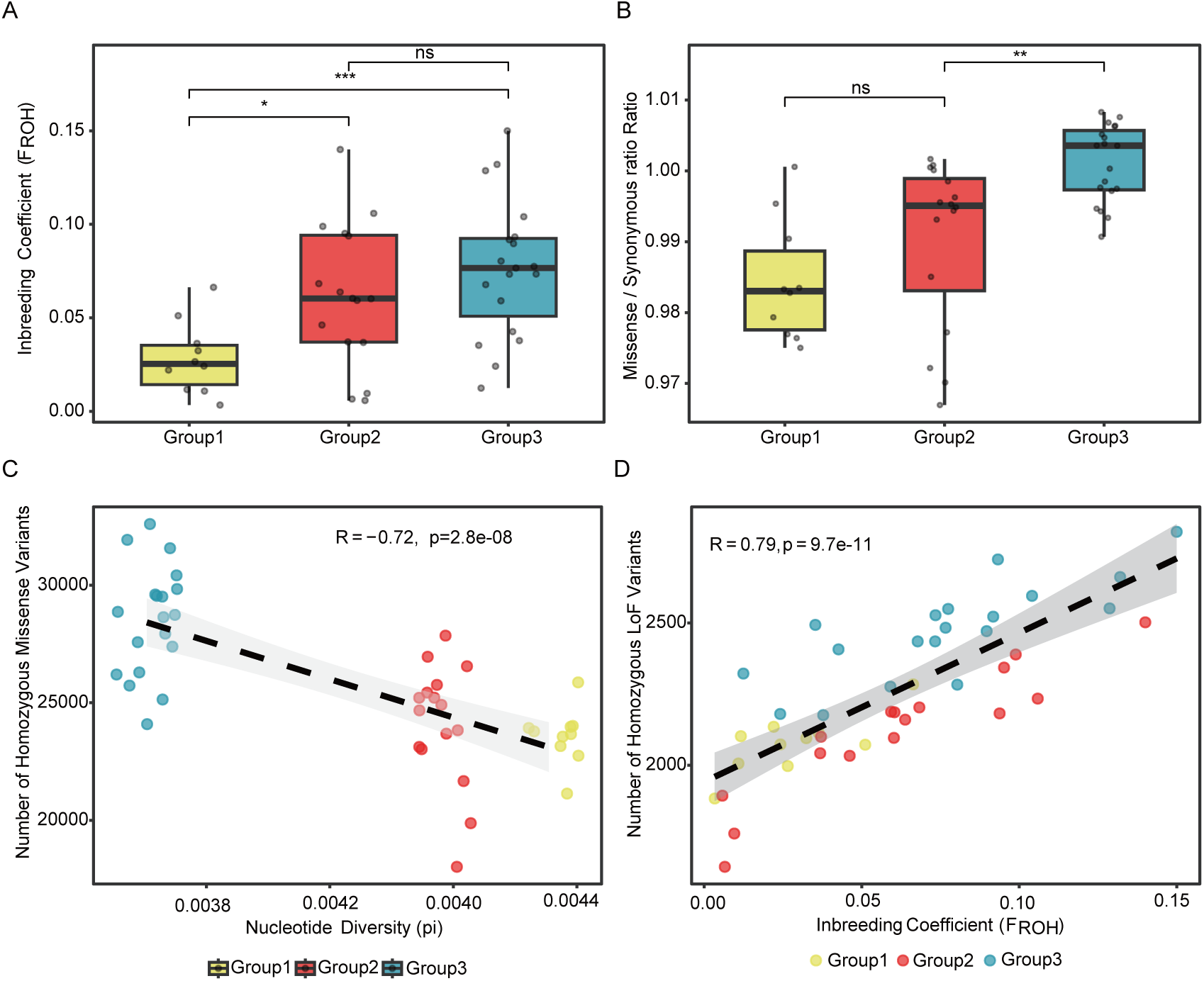
Comparative analysis of genetic diversity, inbreeding, and mutational load. (A). Inbreeding coefficient across populations. Box plots illustrating the distribution of inbreeding coefficients based on runs of homozygosity (ROH). Group 3 (blue) and Group 2 (red) exhibit significantly higher levels of inbreeding compared to Group 1 (yellow), while the outgroup (grey) shows negligible. (B). Purifying selection efficacy measured by dN/dS ratio. Comparison of the ratio of nonsynonymous (missense) to synonymous substitutions among the three genetic groups. Group 3 shows a significantly higher ratio (P < 0.01) than Group 2, suggesting a relaxed efficacy of purifying selection in this lineage. No significant difference (ns) is observed between Group 2 and Group 1. (C). Relationship between genetic diversity and mutational load. Correlation between nucleotide diversity (pi) and the number of homozygous missense variants. A negative correlation is observed, indicating that populations with lower genetic diversity (e.g., Group 3) tend to accumulate a higher burden of deleterious homozygous mutations. (D). Impact of inbreeding on the accumulation of loss-of-function (LoF) variants.Linear regression analysis between the inbreeding coefficient () and the number of homozygous LoF variants. A strong positive correlation suggests that increased inbreeding is a primary driver for the fixation of severe deleterious variants within these earthworm populations.

These pronounced disparities in inbreeding are accompanied by clear contrasts in the efficacy of purifying selection and the accumulation of deleterious genomic variants. Genome-wide functional annotation using SnpEff^17^, polarized by outgroup genotypes, revealed a marked increase in the genetic load within Group 3 (**Table S10**). Specifically, the ratio of missense (deleterious) to synonymous substitutions—a key indicator of selective pressure—is significantly elevated in Group 3 compared with Group 2 (P < 0.01), while no significant difference was observed between Group 1 and Group 2 (**Fig. 3B)**. The higher Mis/Syn Ratio (1.0010) in Group 3 indicates relaxed purifying selection and an increased retention of moderately deleterious mutations.

Further decomposition of mutational counts reveals that the compromised genomic health of Group 3 is driven by the physical accumulation of high-impact variants (**Supplementary Fig. 7**). Consistent with its elevated ROH-based inbreeding coefficients (F_ROH), Group 3 harbors a substantially higher burden of homozygous loss-of-function (LoF) mutations (2468.9 ± 171.5) and homozygous missense variants (28,509.4 ± 2,335.2) compared with Group 1 (LoF: 2076.1 ± 104.2; missense: 23,583.9 ± 1,182.9) (**Supplementary Fig. 7B,C & Table S10**). This pattern suggests that recent population bottlenecks and restricted connectivity have not only increased homozygosity through inbreeding but also reduced the efficacy of purifying selection in removing deleterious variants from the Group 3 lineage.

To further dissect the relationship between genetic diversity, inbreeding, and mutational burden, we examined correlations between genome-wide diversity metrics and deleterious variant accumulation. Across populations, nucleotide diversity (π) shows a strong negative correlation with the burden of homozygous missense variants (R = −0.72, P = 2.8 × 10^−8; **Fig. 3C**), indicating that reduced genetic diversity is associated with an increased susceptibility to mutational load. Consistently, inbreeding coefficients are strongly positively correlated with the accumulation of homozygous loss-of-function (LoF) variants (R = 0.79, P = 9.7 × 10^−11; **Fig. 3D**), highlighting the compounding effects of homozygosity on the exposure and fixation of deleterious alleles.

Collectively, these results demonstrate that broadly shared demographic histories do not necessarily translate into uniform genomic outcomes. Instead, lineage-specific differences in inbreeding intensity and genetic diversity have generated pronounced heterogeneity in genomic integrity among populations of *A. aspergillum*. In particular, elevated inbreeding in Group 3—likely exacerbated by localized habitat fragmentation and complex karst topography in the Guangxi region—has amplified the accumulation and homozygous expression of deleterious variants. These findings highlight inbreeding as a key driver shaping mutational load and genomic health, operating even in the absence of complete geographic isolation and despite broadly synchronous demographic trajectories.

### Adaptive introgression and functional selection underpin environmental resilience

To characterize mitochondrial lineage composition across nuclear-defined groups, we reconstructed a mitochondrial phylogeny based on 13 protein-coding genes and compared it with the nuclear tree using a tanglegram framework (**Fig. 4A**). The mitochondrial genomes clustered into four well-supported clades, hereafter referred to as Mito-G1, Mito-G2, Mito-G3, and Mito-G4, corresponding to the purple, grey, brown, and green backgrounds shown in **Fig. 4A**, respectively. The distribution of mitochondrial haplotypes differed markedly among nuclear genetic groups. Most individuals from nuclear Group 1 (8 out of 10) clustered within Mito-G1, with the remaining two individuals placed in Mito-G2 and Mito-G3, respectively. In contrast, nuclear Group 2 showed a more heterogeneous mitochondrial composition: majority of individuals fell within Mito-G2 and Mito-G3, whereas two individuals (cc3 and X31) were assigned to Mito-G4. Nuclear Group 3 exhibited the highest mitochondrial diversity, with individuals distributed across all four mitochondrial clades, most frequently within Mito-G1 and Mito-G4 (5/19 and 6/19, respectively).

**Figure 4.**
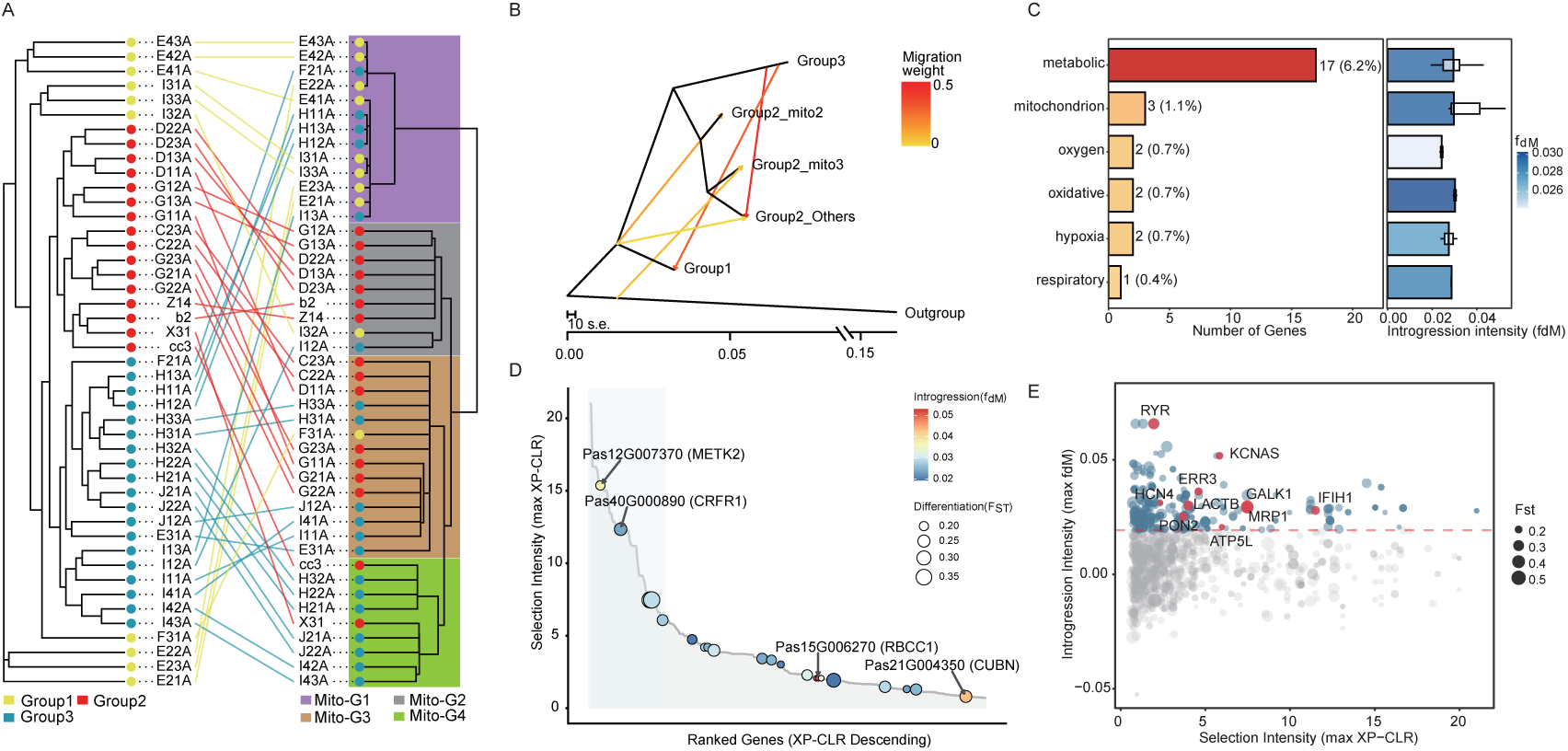
Genome-wide evidence for cytonuclear discordance and adaptive introgression. (A) Tanglegram comparing the nuclear phylogeny inferred from whole-genome SNPs (left) with the mitochondrial phylogeny reconstructed from 13 mitochondrial protein-coding genes (right). Nuclear genetic groups are indicated by colored tips (Group 1, yellow; Group 2, red; Group 3, blue), whereas mitochondrial clades are denoted by background shading (Mito-G1 to Mito-G4). Lines connect identical individuals across trees, revealing pronounced discordance between mitochondrial and nuclear genealogies. (B) Maximum-likelihood population graph inferred using TreeMix with five migration events (m = 5). Arrows represent inferred gene-flow events, with color intensity proportional to migration weight (w). Partitioning Group 2 by mitochondrial affinity reveals introgression from the outgroup into the Group2_mito3 sublineage and from Group 1 into the Group2_mito2 and Group2_other sublineages, uncovering a mosaic genomic architecture shaped by secondary contact. (C) Functional annotation summary of 256 high-confidence candidate genes identified by joint signals of selection (XP-CLR), genetic differentiation (F_ST), and introgression (f_dM). Candidates were classified using an annotation-guided keyword approach based on GO term descriptions. Bar plots show the frequency of genes associated with metabolism-, mitochondrion-, respiration-, and oxygen-related functions, whereas boxplots depict introgression intensity (f_dM) across functional categories. (D) Genome-wide ranking of selection intensity (maximum XP-CLR), with the 21 candidate genes associated with energy metabolism, mitochondrial function, redox processes, and oxygen availability projected onto the distribution. Genes are ordered by decreasing XP-CLR values, while introgression intensity (f_dM) is shown by color gradients. METK2 and CRFR1 rank among the strongest selection signals, whereas RBCC1 and CUBN exhibit the highest introgression intensity within this functional subset. (E) Visualization of selection and introgression signals among functionally annotated candidates. Scatter plot showing selection intensity (XP-CLR, x-axis) versus introgression intensity (f_dM, y-axis) for the 256 candidate genes identified by joint genome-wide signals. Point size reflects genetic differentiation (F_ST). Genes exceeding the predefined thresholds for XP-CLR, f_dM, and F_ST are highlighted in blue. Among these, ten genes identified through cross-species Swiss-Prot annotation as having established roles in mitochondrial respiration, energy metabolism, redox balance, cellular stress responses, or neuromuscular function are highlighted in red.

Although Group 2 forms a cohesive nuclear genetic cluster, its mitochondrial composition is markedly heterogeneous. Among the four high-elevation individuals, only two (Z14 and B2) carry mitochondrial haplotypes concordant with the Group 2 nuclear background, whereas the remaining individuals (X31 and cc3) harbor mitogenomes phylogenetically nested within Group 3 **(Fig. 4A**).

To disentangle these reticulate evolutionary histories, we conducted fine-scale TreeMix^18^ analyses by partitioning Group 2 according to mitochondrial affinity. The optimal number of migration events (m = 5) was determined based on the plateau in the log-likelihood scores (**Supplementary Fig. 8**), and the resulting model exhibited a robust fit with minimal residual variance between most population pairs (**Supplementary Fig. 9**). This refined analysis revealed introgression patterns that were obscured when Group 2 was treated as a single population. Specifically, we detected a significant ancestral genetic contribution from the outgroup into the Group2_mito3 sublineage, alongside a strong migration pulse from the ancestral Group 1 lineage into the Group2_mito2 and Group2_other sublineages (**Fig. 4B**). Together, these results indicate that high-elevation populations of Group 2 have repeatedly incorporated genetic material from multiple evolutionary sources during secondary contact, resulting in a mosaic genomic architecture.

To pinpoint functional loci underlying adaptive introgression, we integrated multiple genome-wide signals—including selection intensity (XP-CLR), genetic differentiation (Fst), and introgression statistics (fdM)—and identified a high-confidence set of 256 candidate genes (**Supplementary Fig. 10 & Table S11 & Methods**). Due to limited functional annotation in annelids, conventional GO enrichment analyses of the 256 candidate genes did not yield statistically significant results. To nevertheless explore their potential functional relevance, we adopted an annotation-guided categorization strategy by screening GO term descriptions for keywords associated with mitochondrial function, energy metabolism, respiration, and oxygen availability (**Table S11**). Using this hypothesis-driven and annotation-guided approach, we identified a subset of 21 candidate genes (from the total set of 256) with GO term descriptions containing keywords related to energy metabolism, mitochondrial function, redox processes, and oxygen availability. Among these, metabolic-related annotations were the most prevalent, representing 17 genes (6.2% of the candidate set), while keywords linked to mitochondrion, respiratory, and hypoxia-associated processes appeared at lower but consistent frequencies (**Fig. 4D, left panel**). Crucially, while metabolism-related genes were more numerous, those specifically associated with the “Oxidative” term exhibited the highest introgression intensity (fdM) among all functional categories, followed by metabolic and mitochondrion (**Fig. 4D, right panel**). This convergence of functional relevance and strong introgression signals highlights the pivotal role of imported mitochondrial alleles in the adaptive evolution of *A. aspergillum*.

To further evaluate whether these energy metabolism–related loci are associated with extreme selection signals, we projected the 21 candidate genes annotated with GO terms related to energy metabolism, mitochondrial function, redox processes, and oxygen availability onto a genome-wide XP-CLR ranking (**Fig. 4D**). Genes were ordered by descending maximum XP-CLR values to provide a relative landscape of selection intensity, while introgression levels were simultaneously visualized using color gradients of fdm. Within this functional subset, Pas12G007370 (METK2) and Pas40G000890 (CRFR1) ranked as the top two genes with the highest XP-CLR values among the 21 metabolism-related candidates, indicating particularly strong signals of selective differentiation. In contrast, Pas15G006270 (RBCC1) and Pas21G004350 (CUBN) exhibited the highest *f_dM_* values within the same subset, suggesting that these loci represent the strongest introgression signals among functionally annotated candidates (**Fig. 4D**). Among these loci, CRFR1 is of particular interest given its established role in hypoxia-related physiological regulation in plateau mammals, where functional variation in this gene contributes to a blunted hypothalamic – pituitary – adrenal (HPA) axis response under chronic low-oxygen conditions. Such modulation is thought to reduce excessive stress responses and enhance long-term physiological stability in hypoxic environments^19^. Although earthworms lack a canonical HPA axis, the identification of CRFR1 as a top-ranked selection candidate suggests that evolutionarily conserved or analogous stress-response pathways may have been targets of adaptive introgression, facilitating metabolic and physiological adjustment to chronic hypoxia.

By contrast, METK2, RBCC1, and CUBN currently lack direct evidence linking them to high-altitude or hypoxia adaptation in the literature. As the enzyme catalyzing the formation of S-adenosylmethionine (SAM), the universal methyl donor, METK2 plays a central role in cellular metabolism and methylation-dependent regulatory processes (UniProt annotation). Its identification as a primary target of positive selection suggests that METK2 may contribute to metabolic reprogramming under chronic environmental stress, potentially by modulating energy utilization and epigenetic regulation in low-oxygen or other highland-associated conditions. RBCC1 (FIP200) is a core component of the autophagy initiation machinery and plays a conserved role in maintaining protein homeostasis and cellular stress tolerance. Functional studies in Drosophila have demonstrated that Fip200 is essential for autophagy and critically regulates growth, metabolic balance, and aging, underscoring its importance in long-term cellular homeostasis. In the context of chronic hypoxia and low-temperature environments, where damaged proteins and organelles accumulate more rapidly, introgression at RBCC1 may facilitate enhanced autophagic capacity and contribute to physiological resilience^20^. whereas CUBN functions in nutrient uptake, including vitamin B12 absorption^21^, which may indirectly influence metabolic efficiency and oxygen-dependent physiological processes. Together, the prominence of these genes in regions of strong selection or introgression suggests their potential involvement in integrated metabolic and stress-tolerance networks, although their precise roles in hypoxic adaptation remain to be functionally validated.

Cross-species Swiss-Prot annotation further substantiated this pattern, revealing that several introgressed loci correspond to 10 genes with well-established roles in mitochondrial respiration, energy metabolism, and cellular stress responses (**Table S11**). Notably, these included LACTB, a mitochondrial protein implicated in maintaining mitochondrial structure^22^ and metabolic homeostasis^23^; ATP5L, a core component of the oxidative phosphorylation machinery (UniProt P68491); and ERR3(ERRγ), a transcriptional regulator known to coordinate mitochondrial biogenesis and energy metabolism^24^.

Additional candidates, such as PON2 and MRP1, are involved in oxidative stress mitigation^25^ and the transport of metabolic by-products^26^, suggesting enhanced cellular resilience under hypoxic and low-temperature conditions. Genes related to ion transport and neuromuscular function, including HCN4, KCNAS, and RYR, were also identified, highlighting potential adjustments in neural excitability and muscle contraction^27–29^ that may be critical for locomotion and physiological performance in cold, low-oxygen environments.

Furthermore, metabolic reprogramming was supported by the presence of GALK1, is a key enzyme in galactose metabolism, catalyzing the phosphorylation of galactose to galactose-1-phosphate^29^, and IFIH1, which has been linked to cellular stress and immune signaling pathways^30^. Together, these genes point to a convergent functional theme in which introgressed loci disproportionately affect mitochondrial function, redox balance, and energy utilization, consistent with adaptive responses to high-altitude and high-latitude environments.

Collectively, these analyses reveal a coherent pattern in which mitochondrial–nuclear discordance, adaptive introgression, and functional selection converge on pathways central to energy metabolism and mitochondrial function in *A. aspergillum*. High-elevation populations, particularly within nuclear Group 2, exhibit mosaic genomic architectures shaped by repeated introgression from divergent lineages. Within these introgressed regions, candidate loci are non-randomly enriched for genes involved in mitochondrial respiration, oxidative metabolism, redox balance, and cellular stress responses, and these loci display disproportionately strong signals of both selection and introgression. Notably, several introgressed genes correspond to well-characterized functional roles: LACTB, implicated in maintaining mitochondrial structure and metabolic homeostasis; ATP5L, a core component of oxidative phosphorylation; ERRγ (ERR3), a transcriptional regulator coordinating mitochondrial biogenesis and energy metabolism; PON2 and MRP1, involved in oxidative stress mitigation and transport of metabolic by-products, respectively; and HCN4, KCNAS, and RYR, which modulate neural excitability and muscle contraction, potentially supporting locomotion and physiological performance under cold and low-oxygen conditions. Additionally, GALK1, a key enzyme catalyzing galactose phosphorylation, and IFIH1, linked to cellular stress and immune signaling pathways, suggest further contributions to metabolic regulation and stress resilience. Together, these findings indicate that adaptive introgression has facilitated functional fine-tuning of mitochondrial and metabolic processes, providing a mechanistic basis for enhanced physiological resilience and environmental tolerance in hypoxic and cold habitats.

## Discussion

### Cytonuclear discordance reveals temporally decoupled evolutionary histories across karst landscapes

Our analyses uncover a pronounced cytonuclear discordance in *Amynthas aspergillum* populations from the Guangxi karst, reconciling deep mitochondrial divergence with shallow nuclear differentiation. Previous phylogeographic studies based on mitochondrial markers documented multiple deeply divergent lineages originating in the Pliocene (∼4.6 Ma) ^6^, suggesting long-term isolation within discrete karst refugia. In contrast, our whole-genome analyses reveal that diversification among nuclear-defined lineages occurred much more recently, during the Late Pleistocene (∼0.14–0.10 Ma), and was accompanied by extensive, asymmetric nuclear introgression.

This disparity points to a marked temporal decoupling between mitochondrial lineage persistence and nuclear genome evolution. While maternally inherited mitochondrial genomes may retain signatures of ancient isolation due to limited dispersal and strong lineage sorting, nuclear genomes have been repeatedly reshaped by episodic gene flow during secondary contact. Such “leaky” isolation challenges classical expectations of strict allopatric divergence in subterranean invertebrates and highlights karst landscapes as dynamic systems that simultaneously preserve deep lineage histories while permitting intermittent genomic connectivity.

### Karst heterogeneity promotes rapid diversification and asymmetric connectivity

Demographic reconstructions using fastsimcoal2 and SMC++ indicate that lineage diversification within Guangxi occurred rapidly during the Late Pleistocene, coinciding with intensified Quaternary climatic oscillations. Repeated cycles of cooling and warming likely amplified ecological heterogeneity across the karst, fragmenting populations into montane refugia, basins, and subterranean habitats that functioned as transient evolutionary islands.

Despite broadly shared early demographic trajectories, connectivity among lineages was highly asymmetric. Group 1 experienced a secondary expansion and maintained higher effective population sizes, whereas Groups 2 and 3 underwent sustained demographic decline. Patterns of gene flow further indicate pronounced lineage-specific connectivity, with substantial introgression from Group 3 into Group 1 but limited exchange involving Group 2. These results suggest that karst topography does not impose uniform barriers to dispersal; instead, it generates uneven permeability, allowing some lineages to repeatedly reconnect while others remain comparatively isolated.

### Cytonuclear discordance exposes hidden introgression within Group 2

Importantly, cytonuclear discordance proved critical for resolving introgression histories that would otherwise remain obscured. When treated as a single population, Group 2 appeared largely isolated, showing minimal evidence of gene flow in genome-wide analyses. However, partitioning Group 2 according to mitochondrial affinity revealed pronounced internal heterogeneity. Fine-scale TreeMix analyses uncovered repeated introgression events into distinct mito-nuclear sublineages, including gene flow from Group 1 and from the outgroup.

These findings demonstrate that coarse population assignments can mask complex reticulate histories, particularly in fragmented landscapes where secondary contact occurs unevenly across space and time. By explicitly leveraging mito-nuclear discordance, we uncover a mosaic genomic architecture within Group 2 that reflects episodic connectivity despite present-day nuclear cohesion.

### Adaptive introgression targets mitochondrial and metabolic pathways

Beyond shaping population structure, gene flow emerged as a key driver of functional adaptation in *A. aspergillum*. Integrating signals of selection (XP-CLR), differentiation (F_ST), and introgression (f_dM), we identified a set of candidate loci disproportionately associated with mitochondrial function, energy metabolism, redox balance, and cellular stress responses. Several introgressed genes—such as LACTB, ATP5L, ERR3, PON2, and MRP1—play well-established roles in mitochondrial respiration, metabolic homeostasis, and oxidative stress mitigation.

Such functions are likely critical for persistence in karst and high-elevation environments, where chronic hypoxia, low temperatures, and fluctuating resource availability impose strong physiological constraints. Rather than relying solely on de novo mutation, *A. aspergillum* appears to have repeatedly acquired pre-adapted alleles from divergent lineages through hybridization. This process of adaptive introgression provides a rapid route to physiological optimization, facilitating ecological expansion across heterogeneous and environmentally challenging habitats.

### Inbreeding, mutational load, and the limits of isolation

Despite the apparent benefits of gene flow, our results also reveal the long-term genomic costs of isolation. Marked differences in inbreeding intensity among lineages translated into pronounced heterogeneity in mutational load. Group 3, in particular, exhibited elevated ROH burdens, reduced nucleotide diversity, relaxed purifying selection, and a substantial accumulation of homozygous loss-of-function and missense variants. Strong correlations between inbreeding coefficients and deleterious variant burden underscore the compounding effects of reduced effective population size and restricted connectivity on genomic integrity.

These findings indicate that broadly shared demographic histories do not necessarily lead to uniform genomic outcomes. Instead, lineage-specific connectivity determines whether populations experience genetic erosion or maintain long-term genomic health. In this context, introgression may play a dual role: facilitating adaptive innovation while also mitigating the accumulation of deleterious variation in expanding or well-connected lineages.

Together, our results establish *A. aspergillum* as a compelling model for understanding how fragmented landscapes simultaneously promote lineage persistence and evolutionary connectivity. By integrating whole-genome data with cytonuclear analyses, we demonstrate how deep mitochondrial divergence can coexist with shallow nuclear differentiation, how adaptive introgression fuels physiological resilience, and how isolation exacerbates mutational load. These insights extend beyond earthworms, offering broader implications for the evolution of subterranean and soil-dwelling organisms inhabiting complex topographies, and highlight the importance of maintaining connectivity to sustain both adaptive potential and genomic integrity.

## Methods

### Sample Collection and Whole-Genome Resequencing

A total of 48 adult *Amynthas aspergillum* individuals were collected from 25 diverse geographic locations across the subtropical landscapes of the Guangxi region, China. To ensure high-quality genomic analysis, total genomic DNA was extracted from the muscle tissue of each individual using a modified phenol-chloroform protocol. Briefly, fresh tissues were flash-frozen and pulverized in liquid nitrogen, followed by homogenization in a lysis buffer [200 mM Tris-HCl (pH 8.0), 20 mM Na_2EDTA, 1% SDS] supplemented with RNase A (2 mg/mL) and Proteinase K (20 mg/mL). After incubation at 56 ° C and subsequent purification steps using phenol/chloroform/isoamyl alcohol (25:24:1), the genomic DNA was precipitated with ammonium acetate and cold ethanol. The integrity and concentration of the extracted DNA were verified using agarose gel electrophoresis and a Qubit Fluorometer (Thermo Fisher Scientific, Waltham, MA, USA).

Whole-genome resequencing was performed using the Illumina HiSeq 2500 platform (Illumina, San Diego, CA, USA) with a 150 bp paired-end strategy. Genomic DNA was sheared into approximately 300 bp fragments using a Covaris focused-ultrasonicator to construct high-quality sequencing libraries. To balance depth and coverage across the dataset, six individuals (Y13, L14 b2, cc3, X31 and Z14) were sequenced at a high coverage of approximately 30X, while the remaining 42 individuals were sequenced at a target depth of 10X. All generated reads were subjected to rigorous quality control to remove low-quality bases and adapter sequences before being aligned to the telomere-to-telomere (T2T) reference genome of *A. aspergillum*.

### Read Mapping and Quality Assessment

Raw sequencing reads were processed to remove adapter sequences and low-quality bases using fastp^31^ (v1.0). The cleaned reads were then aligned to the *A. aspergillum* T2T reference genome using BWA-MEM (v2.2.1)^32^ with default parameters. To ensure high-quality downstream analysis, we performed a comprehensive assessment of the alignment statistics. Mapping rates and the number of mapped reads/bases were quantified using Samtools (v1.22)^33^ stats and MultiQC (v1.12)^34^. To evaluate the uniformity and completeness of genome coverage, we employed mosdepth (v0.3.10)^35^ to calculate the average sequencing depth and the percentage of the genome covered at >1X and >5X depths for each individual. Only samples meeting our stringent quality criteria—specifically a mapping rate >90% were retained for subsequent variant discovery.

### Variant Calling and Quality Control

High-quality reads were aligned to the *A. aspergillum* T2T reference genome using BWA-MEM2 (v2.2.1)^32^. Alignments were coordinate-sorted via Samtools (v1.22)^33^ and processed with GATK (v 4.6.2.0)^36^. PCR duplicates were identified and marked using MarkDuplicatesSpark. Individual-level variant discovery was performed with GATK HaplotypeCaller in–ERC GVCF mode to record genotype likelihoods across all sites. Individual GVCFs were consolidated into a GenomicsDB, and joint genotyping was conducted using GenotypeGVCFs to generate a population-level VCF comprising 48 individuals. We implemented a multi-stage filtering pipeline to ensure a high-confidence SNP set. Initial quality control was executed using BCFtools (v1.22) ^33^, retaining only bi-allelic SNPs. We applied stringent GATK hard-filtering (QUAL > 30, MQ > 40, FS < 60, SOR < 3) and variants overlapping annotated transposable elements (TEs) in the T2T genome were systematically masked. At the genotype level, calls with a read depth (DP) < 5 were treated as missing data. Final site-level filtering excluded variants with a missing rate >10% or a minor allele frequency (MAF) < 0.05.

### Species Identification and Outgroup Selection

We implemented a dual-level identification strategy integrating organellar and nuclear genomic data. First, complete mitochondrial genomes were de novo assembled for all individuals using GetOrganelle (v1.7.5)^37^. The assembled mitogenomes were combined with the published mitochondrial genome of *Amynthas robustus* (GenBank accession LC726556.1) for phylogenetic analyses to assess mitochondrial lineage clustering.

### Population Genetic Structure and Phylogenetic Inference

Population structure and phylogenetic relationships were characterized using a filtered SNP subset pruned for linkage disequilibrium (LD). LD pruning was executed in PLINK (v1.90)^38^ with a sliding window approach (--indep-pairwise 50 10 0.2). Ancestral proportions were estimated for the complete dataset of 48 individuals using ADMIXTURE (v1.3.0)^11^ across a range of ancestral clusters (K = 2 to 7) (**Supplementary Fig. 4**). Principal Component Analysis (PCA) was performed on the 45 *A. aspergillum* individuals (excluding the outgroup: Y13, L14 and B28) using the --pca function in PLINK, with the first ten principal components calculated to define the genetic clusters. Phylogenetic relationships were resolved through maximum-likelihood (ML) inference. The filtered VCF was converted to a phylip alignment using vcf2phylip^39^, with the three *A. robustus*-like individuals designated as the outgroup. The ML tree was constructed in IQ-TREE (v2.2.0)^40^ employing the GTR+G substitution model and the -fast option. Nodal support was assessed using 1,000 replicates of the Ultrafast Bootstrap (UFBoot) method.

### Estimation of Individual Heterozygosity and Genetic Diversity

To assess the level of genetic variation within the sampled populations, we quantified individual heterozygosity across the final set of about 10.8 million high-quality SNPs. For each of the 45 individuals, the number of heterozygous (nHet) and homozygous (nHom) genotypes was enumerated to calculate the genome-wide heterozygosity rate (H = nHet / (nHet + nHom)). These statistics were summarized for each of the identified genetic groups. Nucleotide diversity (pi) and linkage disequilibrium (LD) decay were further estimated to characterize the genetic landscape of each group. Specifically, pi was calculated using a sliding-window approach (window size 50 kb, step size 10 kb) via VCFtools (v0.1.18)^41^. To evaluate the rate of LD decay, the squared correlation coefficient (r^2) between pairs of SNPs was calculated using PopLDdecay (v3.43)^42^ with a maximum distance of 100 kb. These metrics provided the quantitative basis for evaluating the impacts of historical bottlenecks and localized inbreeding on the genomic integrity of *A. aspergillum*.

### Divergence Time Estimation and Demographic History

Single-copy orthologous genes were identified across four representative species using OrthoFinder (v2.5.5)^43^. Divergence times were estimated based on a concatenated dataset of 585 orthologs using MCMCtree in PAML (v4.9). The analysis employed an independent rates clock model (clock = 2) and the HKY85 substitution model. Temporal constraints were implemented using fossil-calibrated nodes derived from the TimeTree database (http://www.timetree.org/). Divergence times were estimated using the MCMCtree program in PAML^12^. The MCMC chain was executed for 200,000 iterations, with the initial 20,000 iterations discarded as burn-in. Post-burn-in trees were sampled every 10 iterations until all parameters reached an effective sample size (ESS) > 200. Subsequently, long-term effective population size (Ne) fluctuations were reconstructed using SMC++ (v1.15.4) ^15^, integrating site frequency spectrum (SFS) and linkage disequilibrium (LD) data. Trajectories were estimated independently for each of the four genetic groups using a mutation rate (mu) of 1.18 * 10^-9 and a generation time (g) of 0.6 years.

### Demographic Modeling and Gene Flow

Population splitting and migration events were characterized using TreeMix (v1.13) ^18^ and fastsimcoal2 (v2.8) ^16^. TreeMix modeled migration (m = 0–5) using a block bootstrap (k = 10,000 SNPs), with three A. robustus-like individuals as the outgroup. Optimal edges were determined by variance explained and bootstrap stability. Demographic parameters were further quantified via SFS-based coalescent simulations in fastsimcoal2. The finalized model incorporated three sequential divergence events and bi-directional migration between Group 1 and Group 3. Parameter estimation was based on 100 independent runs (100,000 simulations and 40 ECM cycles per run), with the best-fit model identified by the maximum observed likelihood.

### Variant Functional Annotation and Mutational Load Assessment

Functional consequences of all identified variants were predicted using snpEff (v5.2) ^17^ based on the *A. aspergillum* T2T reference genome and its associated gene models. Variants were categorized into four functional classes: synonymous, missense (deleterious), and loss-of-function (LoF). LoF variants were strictly defined as those leading to premature stop-gain, frameshift, or the disruption of essential splice sites. To evaluate the genetic load across the 45 individuals, we calculated three complementary metrics: (i) the total number of derived alleles per individual, (ii) the number of heterozygous variants, and (iii) the number of homozygous variants for each functional category. The efficacy of purifying selection was further quantified by calculating the ratio of missense to synonymous substitutions for each population.

### Characterization of Runs of Homozygosity (ROH)

Individual inbreeding coefficients R_OH_ were estimated by identifying runs of homozygosity (ROH) using PLINK (v1.90). We implemented a sliding window approach with the following stringent parameters to ensure high-confidence ROH detection: a minimum window size of 50 SNPs (--homozyg-window-snp 50), a minimum ROH length of 300 kb (--homozyg-kb 300), a required density of one SNP per 50 kb (--homozyg-density 50), and a maximum of one heterozygous call allowed per window (--homozyg-window-het 1). Detected ROHs were classified into three length categories: short (0.3–1.0 Mb), medium (1.0–5.0 Mb), and long (>5.0 Mb). F_ROH was calculated as the total length of all ROHs divided by the total length of the assembled genome (758.86 Mb).

### Correlation Analysis and Statistical Inference

Statistical comparisons of mutational load and inbreeding coefficients between genetic groups were performed using the Wilcoxon rank-sum test. The relationship between genome-wide nucleotide diversity (pi), inbreeding (F_ROH), and the accumulation of deleterious variants was evaluated via Pearson correlation analysis. Linear regression was employed to model the impact of inbreeding on the fixation of homozygous LoF variants. All statistical analyses and visualizations were executed in R (v4.5) using the ggplot2, ggpubr, and patchwork packages.

### Genome-wide Selection Scans

Genomic regions under positive selection were identified through a cross-population composite likelihood ratio (XP-CLR v1.5)^44^ analysis and fixation index (F_ST) calculations. Genetic differentiation between the target group (Group2_mito3) and the reference group (Group2_Others) was quantified using VCFtools (v0.1.18)^41^, employing a sliding window of 50 kb with a 10 kb step size. XP-CLR was implemented with a window size of 50 kb and a 10 kb step size, enforcing a minimum of 10 SNPs and a maximum of 600 SNPs per window. Linkage disequilibrium (LD) was weighted with a threshold of 0.95. Candidate selective sweeps were defined by the intersection of the top 1% empirical distributions of both F_ST and XP-CLR scores.To identify specific functional targets of selection, candidate windows were intersected with mitochondrial-related gene models using bedtools map (v2.29.1)^45^. For each gene, the maximum F_ST and XP-CLR values within its genomic coordinates were extracted to characterize its selective significance.

### Localized Introgression and Adaptive Gene Flow

Localized genomic introgression was detected using the f_dM statistic, which is specifically suited for evaluating gene flow in small genomic windows. This analysis was performed via Dsuite (v0.5r58)^46^ using the Dinvestigate module. Allele sharing patterns were tested among a triadic configuration: Group2_Others (P1), Group2_mito3 (P2), and the outgroup (O). Calculations were executed in sliding windows of 50 informative SNPs with a step size of 25 SNPs. Statistical significance was assessed using Z-scores derived from a block-jackknife procedure. Genomic regions exhibiting f_dM values in the extreme upper tail (top 5th percentile) of the genome-wide distribution were prioritized as regions of putative adaptive introgression.

## Data availability

All data generated for this study and used in analyses are available in Supplementary Tables.

## Code availability

Data handling and analyses were implemented using standard methods, software tools and code functions detailed in Methods.

## Acknowledgements

This work was supported by the Guangxi Key Research and Development Program, the Guangxi Natural Science Foundation (Grant No. 2025GXNSFAA202689), and the Forestry Science and Technology Project of the Forestry Bureau of Guangxi Zhuang Autonomous Region (Grant No. 2023GXZCLK57). The authors also acknowledge institutional support from their affiliated institutions. The funding agencies had no role in study design, data analysis, interpretation, or manuscript preparation.

## Ethics declarations

