## Supplementary_Figures for "Genomic insights into rapid diversification and adaptive introgression in the earthworm *Amynthas aspergillum*"

**Supplementary Fig. 1**

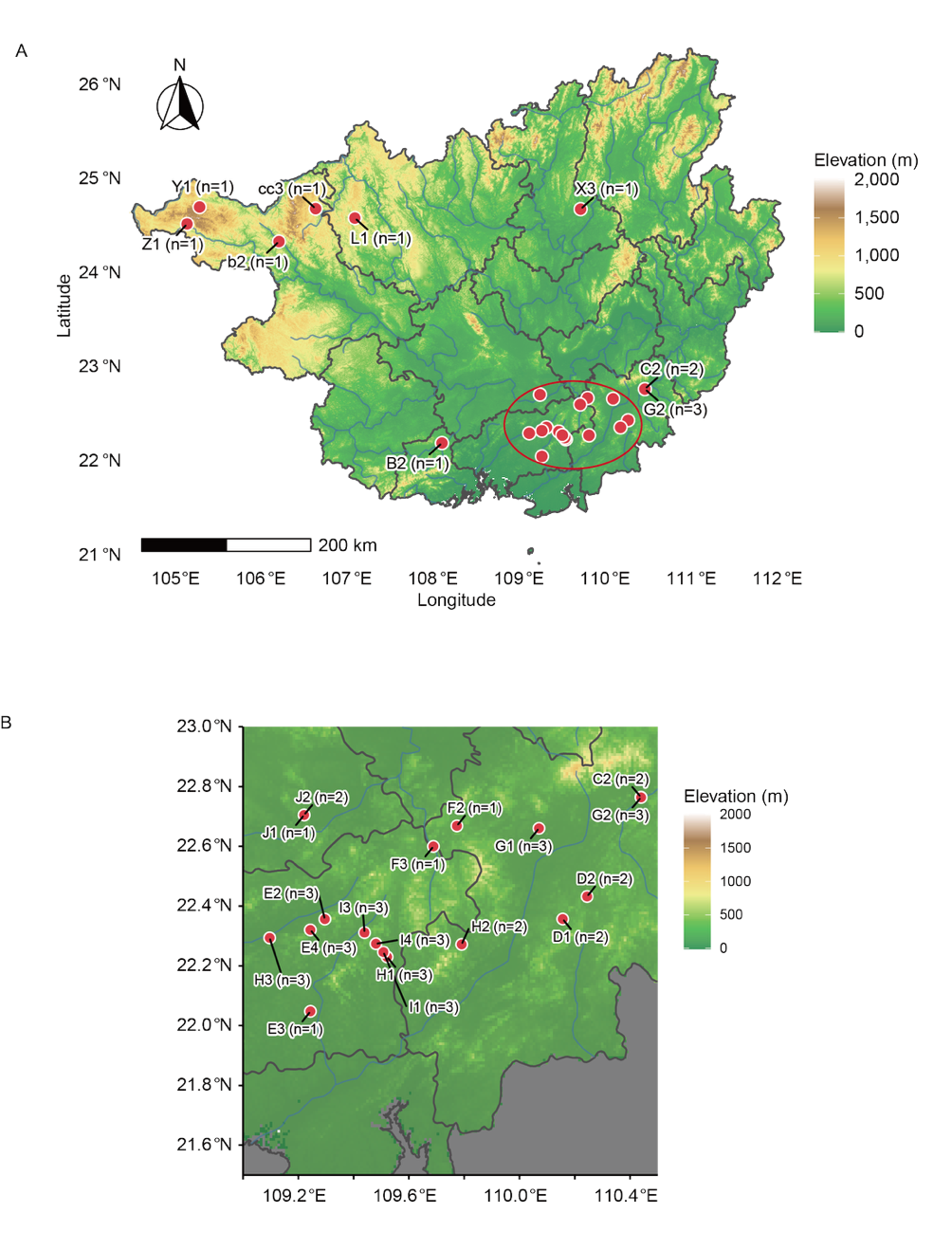

**Supplementary Fig. 1 | Geographic sampling of Amynthas aspergillum across Guangxi, China.** (A) Map showing the geographic locations of 25 sampling sites across the Guangxi region, China, from which 48 Amynthas aspergillum individuals were collected for high-depth whole-genome resequencing (WGS). Sampling sites are indicated by red points, and the numbers in brackets denote the number of individuals sampled at each site. (B) Local zoom-in view of the sampling sites shown in panel (A), highlighted by red circles.

**Supplementary Fig. 2**

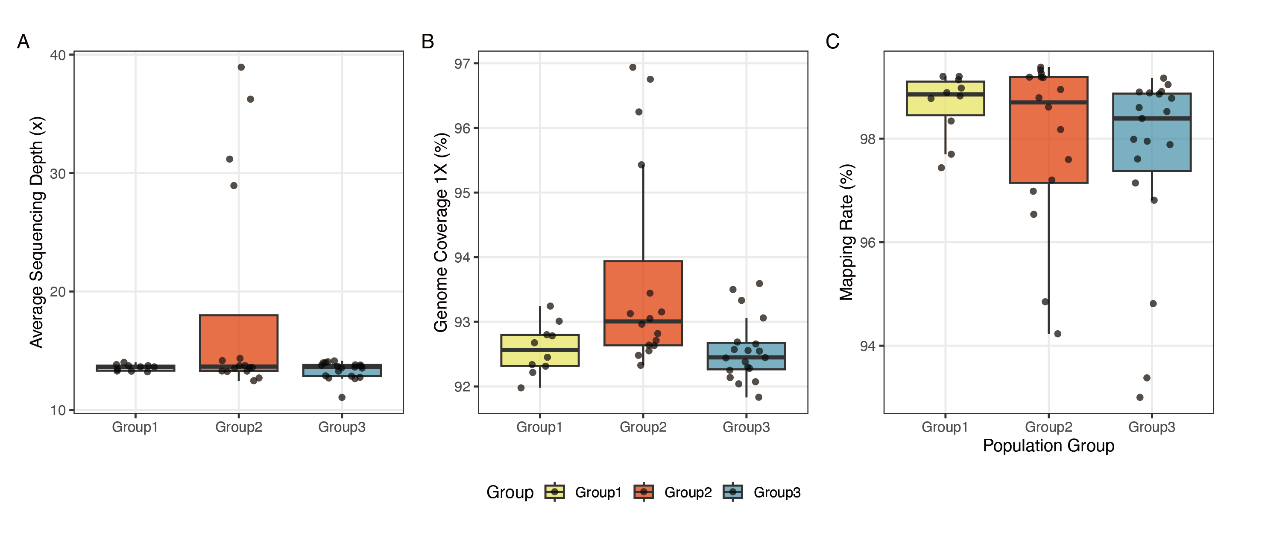

**Supplementary Fig. 2 | Quality control metrics of whole-genome sequencing and read mapping. (**A) Average sequencing depth. Distribution of mean sequencing coverage per individual in each group. Median sequencing depth is comparable among groups. (B) Proportion of the reference genome covered by at least one read for each individual. All groups exhibit high genome coverage, with median values exceeding 92%. (C) Proportion of raw reads successfully aligned to the reference genome. The median mapping rate across all samples is 98.56%.

**Supplementary Fig.** 3

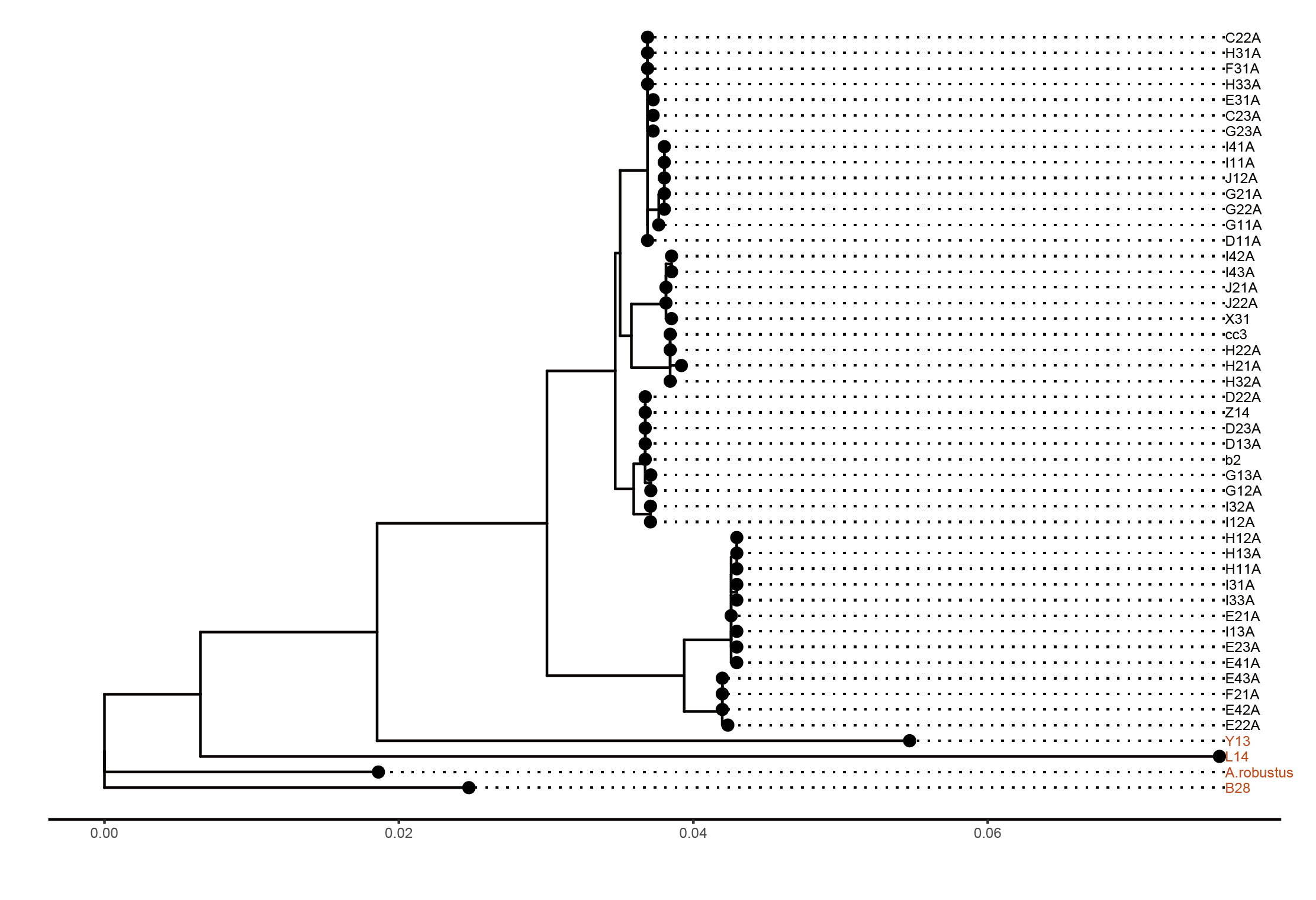

**Supplementary Fig. 3 | Maximum-likelihood phylogeny based on mitochondrial genomes.** Maximum-likelihood (ML) phylogenetic tree inferred from complete mitochondrial genomes of Amynthas aspergillum individuals and related outgroup taxa. The tree depicts maternal lineage relationships among all sampled individuals. Three individuals (B28, L14, and Y13), highlighted in red, cluster closely with the outgroup species A. robustus. The remaining individuals form a distinct and well-supported clade separated from the A. robustus-associated lineage.

**Supplementary Fig. 4**
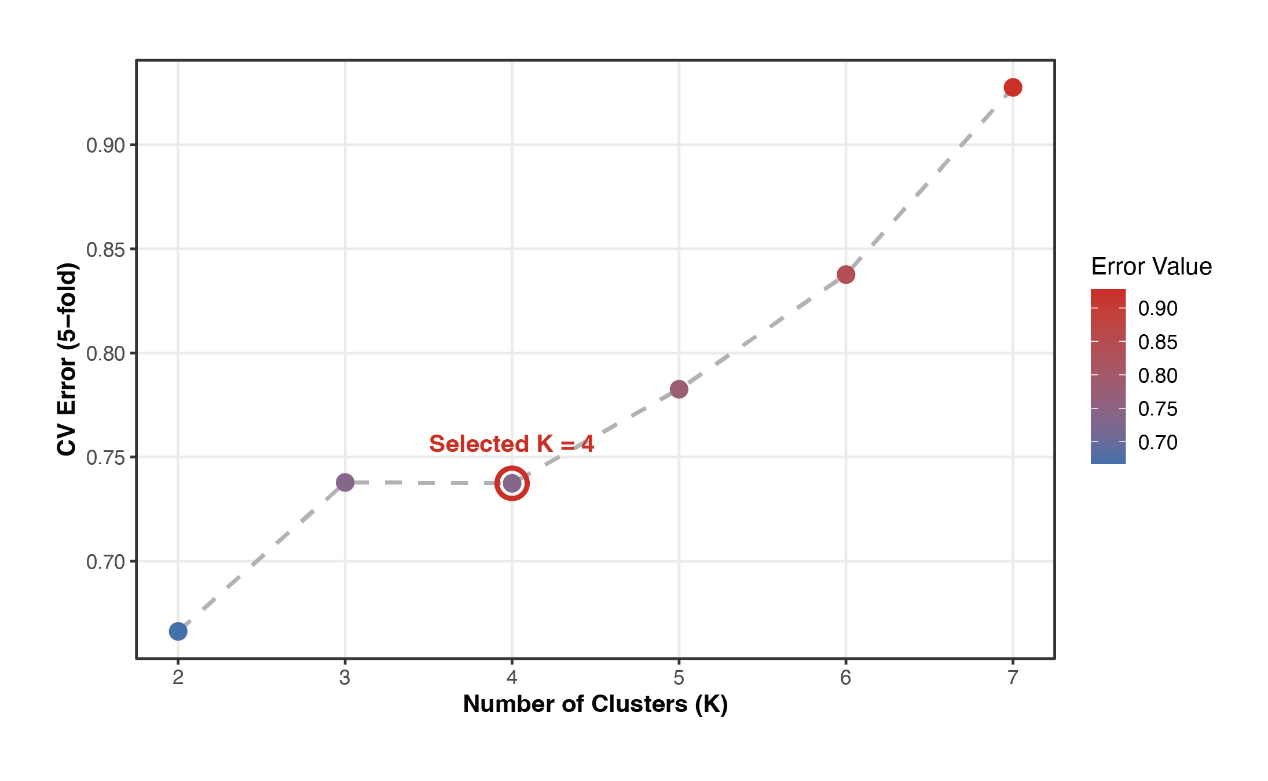

**Supplementary Fig. 4 | Cross-validation of ADMIXTURE models. Cross-validation (CV) error scores for ADMIXTURE models with ancestral cluster numbers ranging from K = 2 to K = 7. Five-fold cross-validation was performed for each K value to assess model fit and identify the optimal number of ancestral populations. The lowest CV error was observed at K = 2; however, this model did not capture the finer-scale genetic structure evident in the principal component analysis (PCA). Based on biological interpretability and consistency with major PCA axes, K = 4 was selected for downstream nuclear genomic analyses and is highlighted by a red circle.**

**Supplementary Fig. 5**

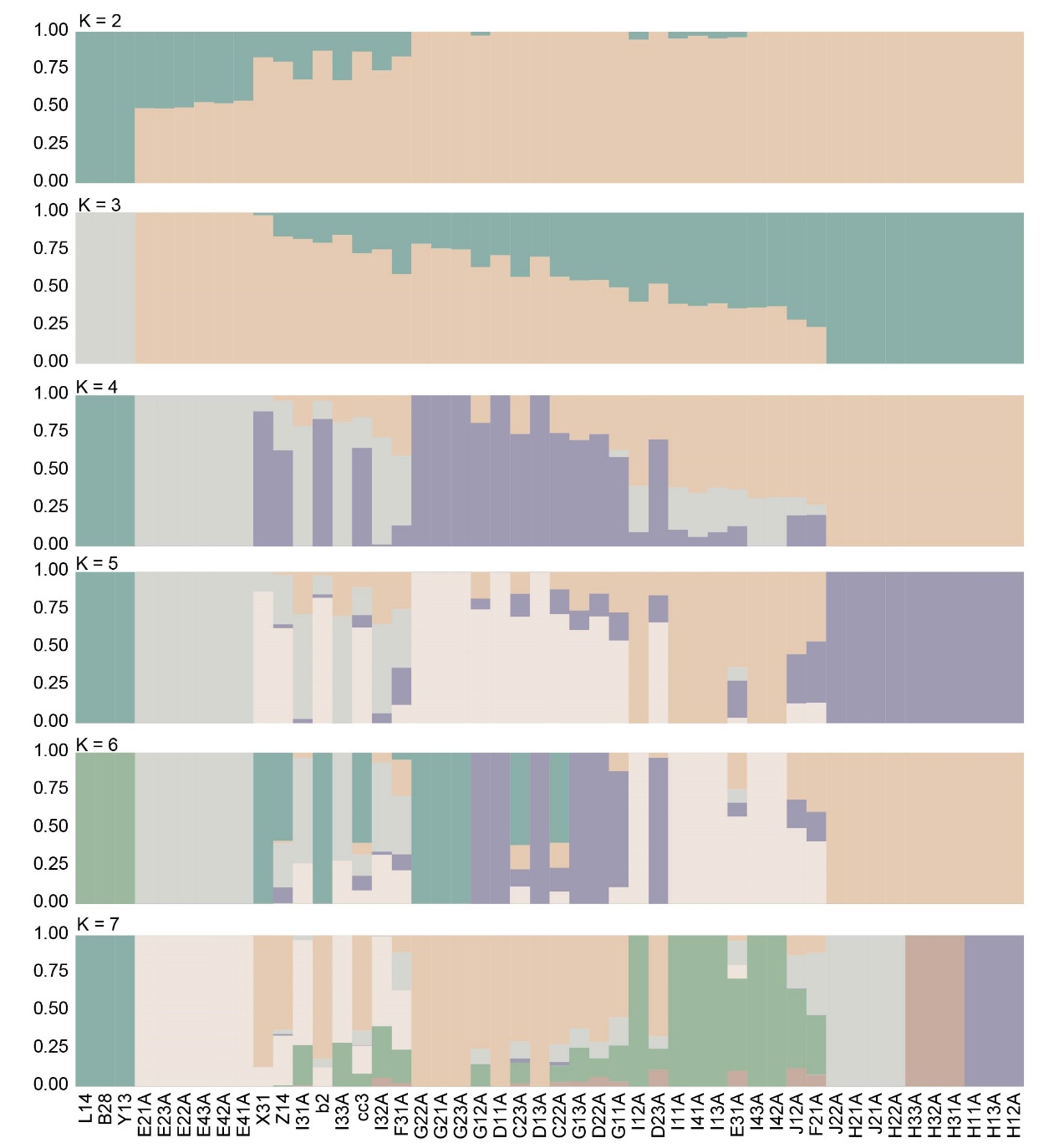

**Supplementary Fig. 5 | Population structure inferred by ADMIXTURE analyses. Ancestry proportions of all individuals estimated using ADMIXTURE for K values ranging from 2 to 7. Each vertical bar represents one individual, and colors indicate the proportion of ancestry assigned to inferred ancestral clusters. At K = 4, population structure is resolved with greater clarity, distinguishing the three primary genetic groups as well as the outgroup individuals (B28, L14, and Y13). Lower K values (K = 2 and 3) do not capture finer-scale genetic structure observed in principal component analysis (PCA), whereas higher K values (K ≥ 5) introduce additional substructure without clear geographic or phylogenetic support. Across all K values, individuals B28, L14, and Y13 consistently display distinct ancestry profiles, supporting their treatment as an outgroup in subsequent nuclear genomic analyses.**

**Supplementary Fig. 6**

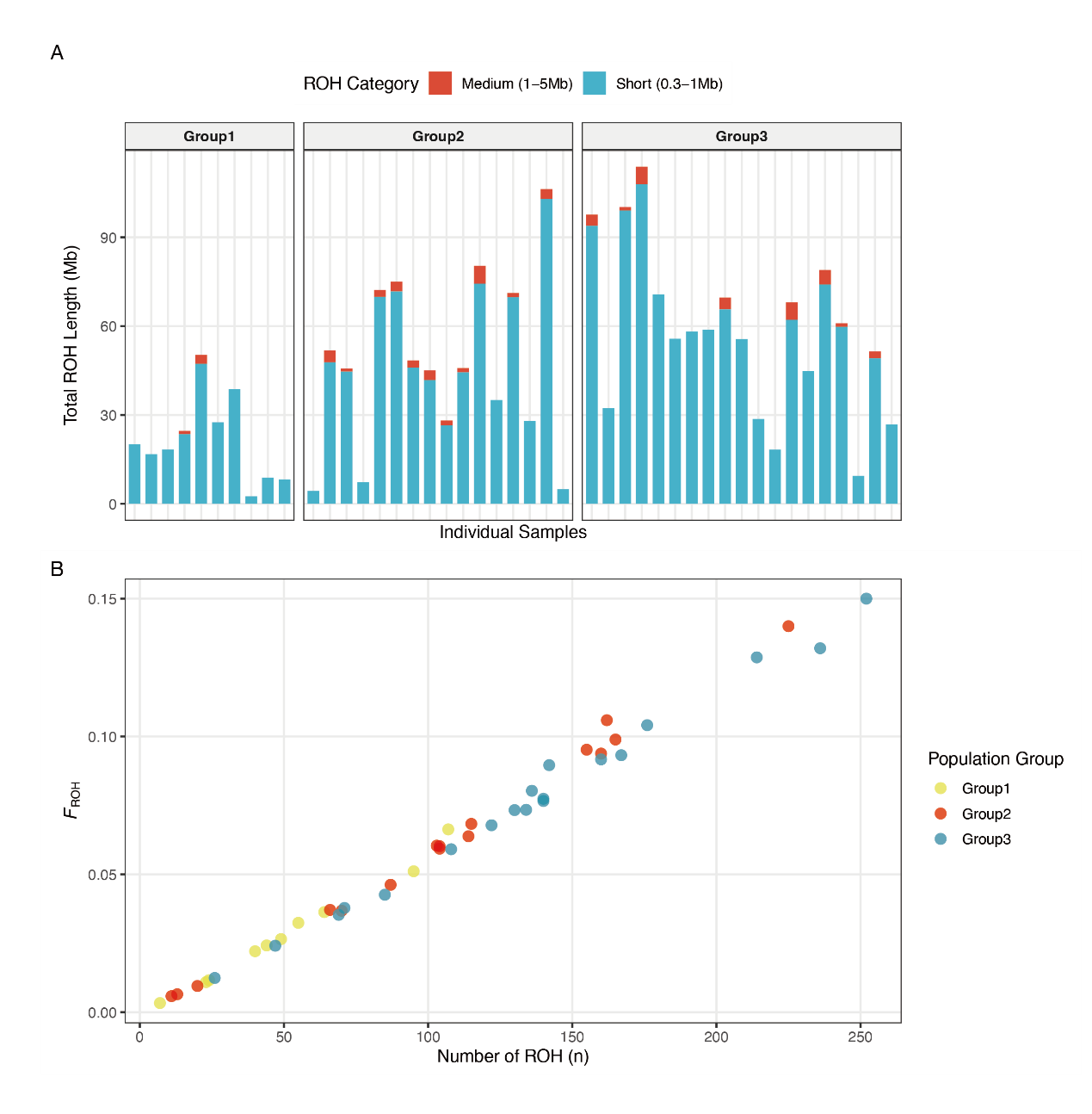

**Supplementary Fig. 6 | Genomic homozygosity patterns and inbreeding levels in A. aspergillum populations.** (A) Total length of ROH per individual, categorized by segment size: short (0.3–1 Mb, light blue) and medium (1–5 Mb, dark blue). Individuals are ordered by population assignment (Group 1, Group 2, and Group 3). (B) Correlation between ROH Count and F_ROH. Scatter plot illustrating the relationship between the total number of ROH segments (N_ROH) and the ROH-based inbreeding coefficient (F_ROH). Points are color-coded by population group: Group 1 (yellow), Group 2 (red), and Group 3 (cyan).

**Supplementary Fig. 7**

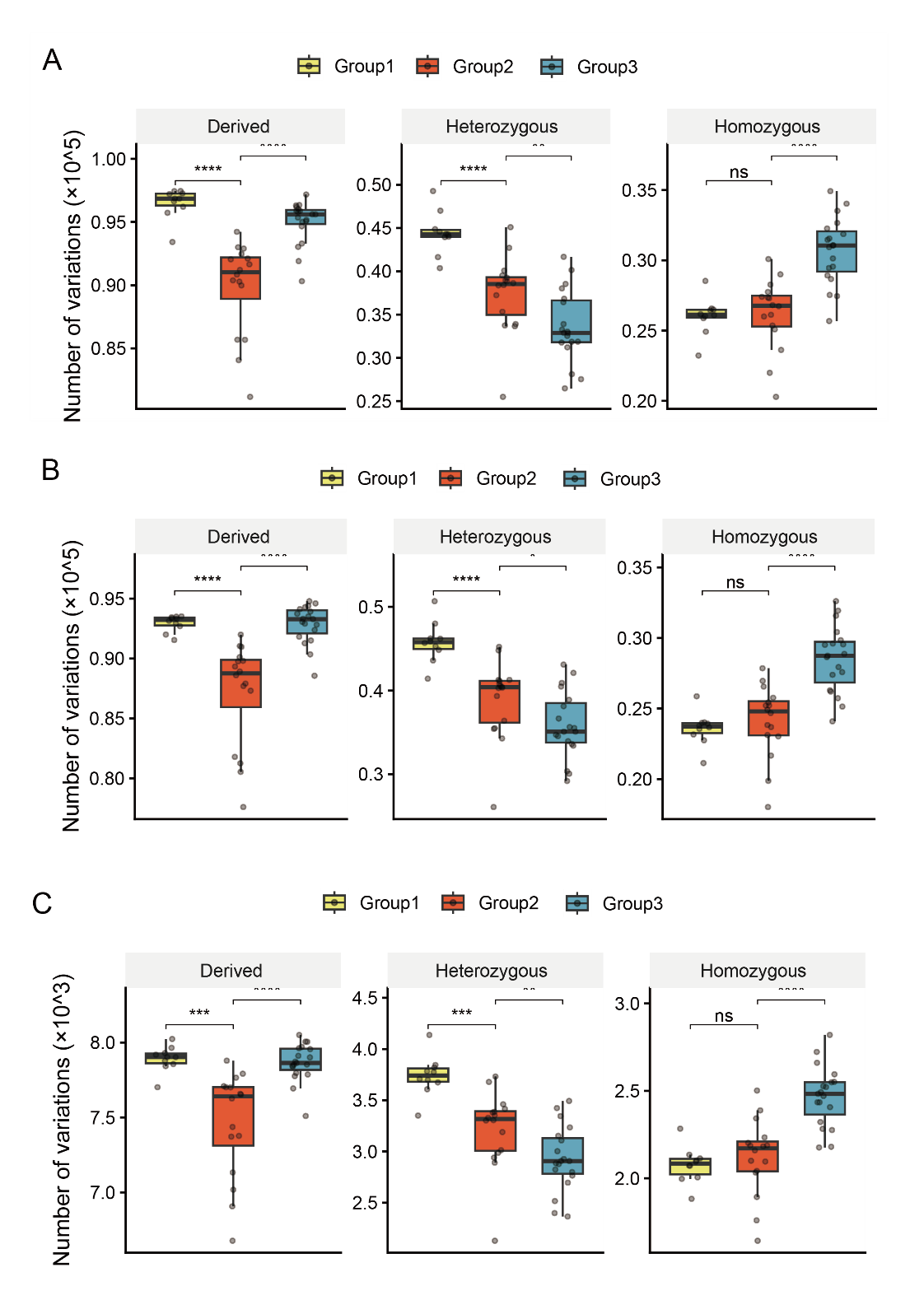

**Supplementary Fig. 7 | Genetic load and mutational burden across population groups.** Comparison of the accumulation and distribution of functional genomic variants among Amynthas aspergillum lineages. Variants were polarized using outgroup genotypes to infer derived alleles. (A) Synonymous variants. Numbers of derived synonymous substitutions in heterozygous and homozygous states for each population group. (B) Missense (deleterious) variants. Distribution of derived missense variants across the three groups, partitioned by zygosity. Group 3 displays an elevated number of homozygous deleterious variants relative to the other groups. (C) Loss-of-function (LoF) variants. Burden of high-impact derived variants, including stop-gain and frameshift mutations. Group 3 shows a higher count of homozygous LoF variants compared with Groups 1 and 2.

Statistical significance was assessed using the Wilcoxon rank-sum test (ns, P > 0.05; ***, P < 0.001; ****, P < 0.0001). Boxes indicate the interquartile range (IQR), and points represent individual samples. Colors denote population groups: Group 1 (yellow), Group 2 (red), and Group 3 (cyan).

**Supplementary Fig. 8**

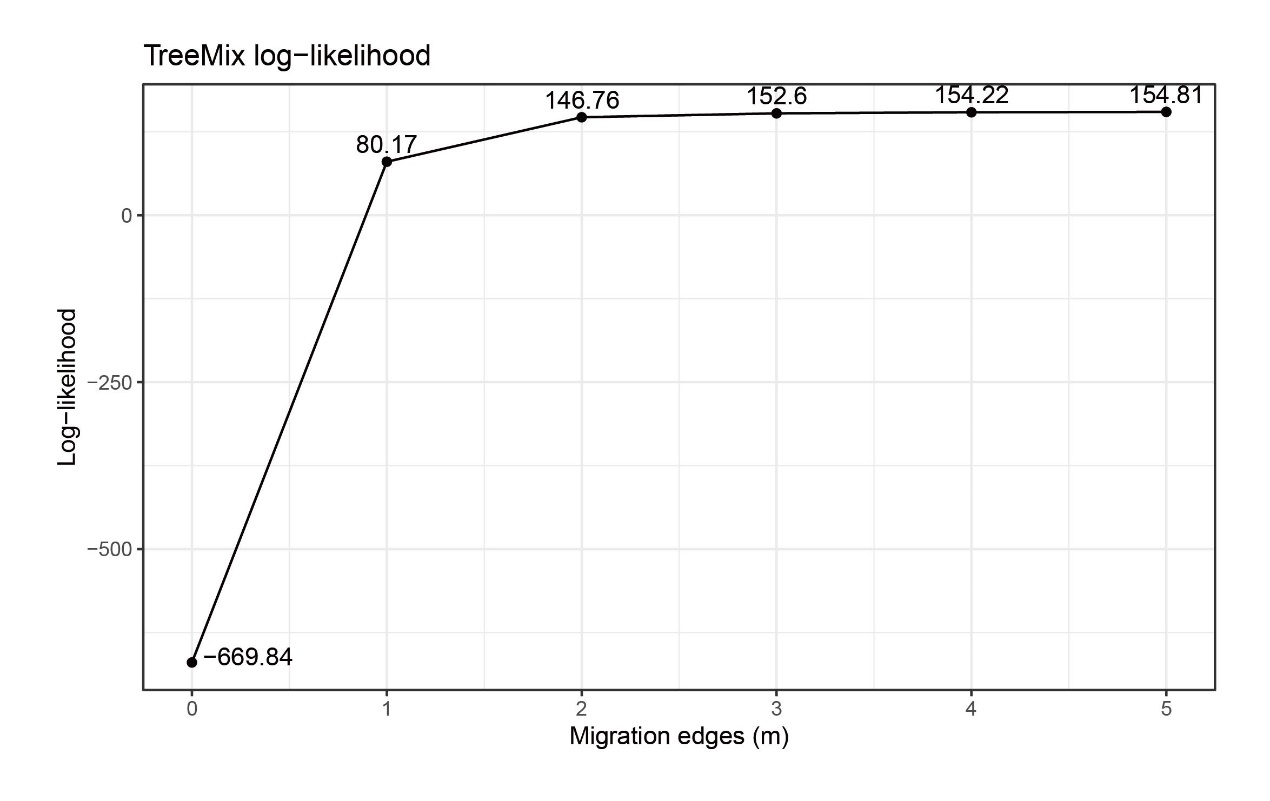

**Supplementary Fig. 8 | Log-likelihood scores for TreeMix models with varying numbers of migration events.** Optimization of the migration model. The line plot shows how log-likelihood changes as the number of migration edges (m) increases from 0 to 5. The scores show a sharp increase at m = 1 and reach a plateau starting at m = 4, supporting the choice of m = 5 to capture the complex reticulate history while avoiding overfitting.

**Supplementary Fig. 9**

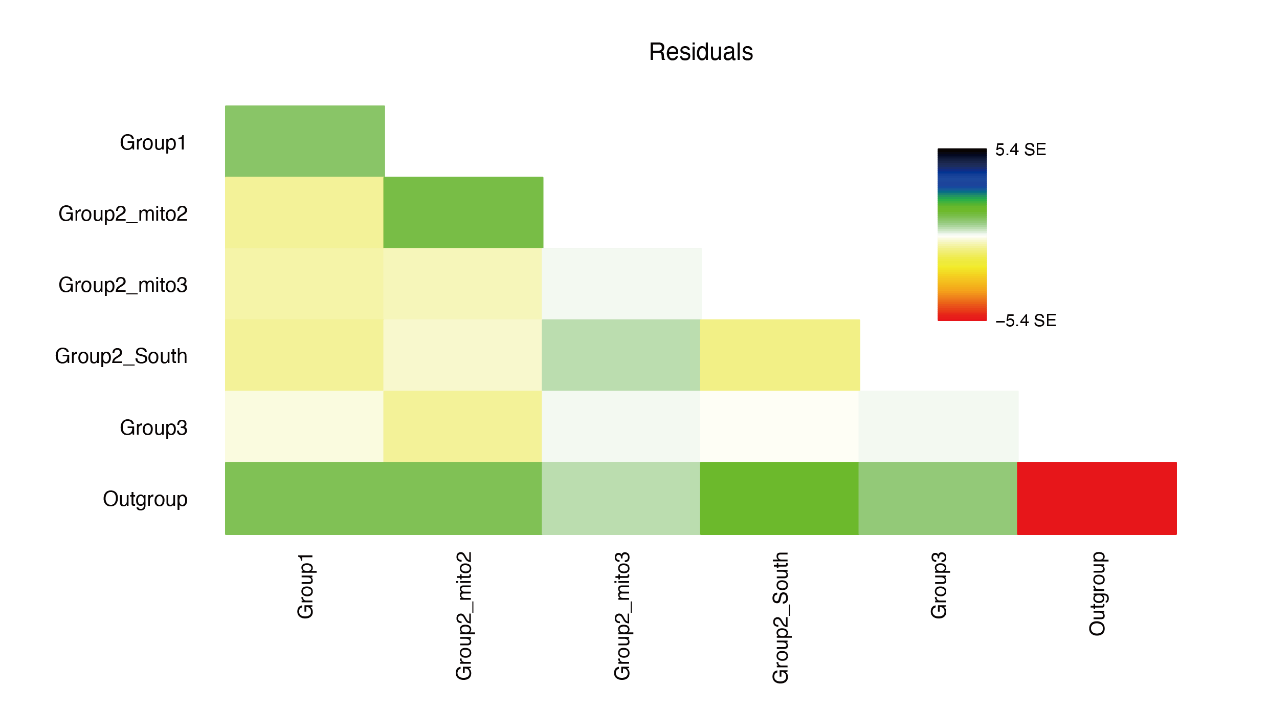

**Supplementary Fig. 9 | Residual fit of the TreeMix model with five migration events.** Residual covariance matrix illustrating the fit of the population tree. The residuals represent the differences between the observed and predicted genetic covariances between population pairs. Most residuals are close to zero (shown as white or light-colored cells), indicating that the model with five migration events accurately captures the major genetic relationships and introgression patterns across lineages. Positive residuals between specific groups suggest potential additional gene flow not represented in the model.

**Supplementary Fig. 10**

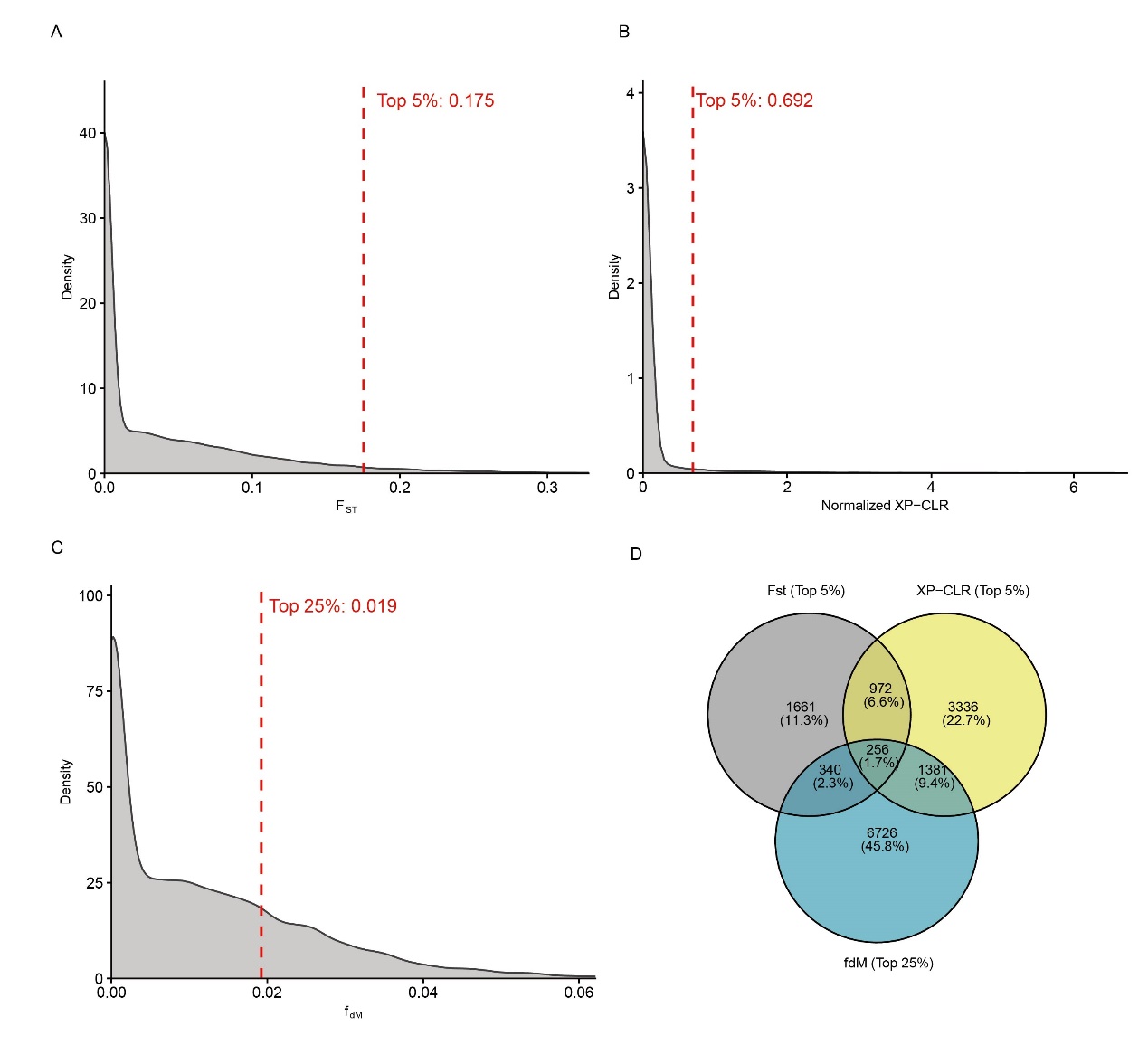

**Supplementary Figure 10. Identification of candidate genes for adaptive introgression.** Integration of genome-wide selection and introgression signals to pinpoint functional loci. (A–C) Genome-wide score distributions: Density plots showing the distributions of (A) genetic differentiation (F_ST), (B) selection intensity (normalized XP-CLR), and (C) introgression statistics (f_dM). Dashed red lines indicate the thresholds used for filtering: the top 5th percentile for F_ST (0.175) and XP-CLR (0.692), and the top 25th percentile for f_dM (0.019). Negative values in F_ST and f_dM were truncated to zero for visualization to minimize statistical noise. (D) Intersection of candidate loci: Venn diagram showing the overlap among genes identified by the three independent metrics. A high-confidence set of 256 candidate genes (central intersection) was identified, representing loci with simultaneous signals of extreme differentiation, strong positive selection, and significant ancestral introgression.
