## Supplemental Data 1 for "Genomic insights into rapid diversification and adaptive introgression in the earthworm *Amynthas aspergillum*"

Table S1. Detailed information on the 48 samples used for whole-genome resequencing.

| SampleID | Locality | Latitude (N) | Longitude (E) |
|---|---|---|---|
| **E21A** | E2 | 22°21'23″N | 109°17'45″E |
| **E22A** | E2 | 22°21'23″N | 109°17'45″E |
| **E23A** | E2 | 22°21'23″N | 109°17'45″E |
| **E41A** | E4 | 22°19'13″N | 109°14'37″E |
| **E42A** | E4 | 22°19'13″N | 109°14'37″E |
| **E43A** | E4 | 22°19'13″N | 109°14'37″E |
| **F21A** | F2 | 22°40'06″N | 109°46'25″E |
| **H11A** | H1 | 22°13'48″N | 109°31'16″E |
| **H12A** | H1 | 22°13'48″N | 109°31'16″E |
| **H13A** | H1 | 22°13'48″N | 109°31'16″E |
| **I13A** | I1 | 22°14'45″N | 109°30'28″E |
| **I31A** | I3 | 22°18'40″N | 109°26'22″E |
| **I33A** | I3 | 22°18'40″N | 109°26'22″E |
| **D22A** | D2 | 22°25'55″N | 110°14'40″E |
| **D23A** | D2 | 22°25'55″N | 110°14'40″E |
| **G12A** | G1 | 22°39'37″N | 110°04'13″E |
| **G13A** | G1 | 22°39'37″N | 110°04'13″E |
| **I12A** | I1 | 22°14'45″N | 109°30'28″E |
| **I32A** | I3 | 22°18'40″N | 109°26'22″E |
| **D13A** | D1 | 22°21'24″N | 110°09'23″E |
| **H21A** | H2 | 22°16'18″N | 109°47'27″E |
| **H22A** | H2 | 22°16'18″N | 109°47'27″E |
| **H32A** | H3 | 22°17'37″N | 109°05'46″E |
| **J22A** | J2 | 22°42'17″N | 109°13'15″E |
| **J21A** | J2 | 22°42'17″N | 109°13'15″E |
| **I42A** | I4 | 22°16'26″N | 109°28'51″E |
| **I43A** | I4 | 22°16'26″N | 109°28'51″E |
| **C22A** | C2 | 22°45'52″N | 110°26'21″E |
| **C23A** | C2 | 22°45'52″N | 110°26'21″E |
| **D11A** | D1 | 22°21'24″N | 110°09'23″E |
| **E31A** | E3 | 22°02'52″N | 109°14'38″E |
| **F31A** | F3 | 22°35'59″N | 109°41'19″E |
| **G11A** | G1 | 22°39'37″N | 110°04'13″E |
| **G21A** | G2 | 22°45'49″N | 110°26'23″E |
| **G22A** | G2 | 22°45'49″N | 110°26'23″E |
| **G23A** | G2 | 22°45'49″N | 110°26'23″E |
| **H31A** | H3 | 22°17'37″N | 109°05'46″E |
| **H33A** | H3 | 22°17'37″N | 109°05'46″E |
| **I11A** | I1 | 22°14'45″N | 109°30'28″E |
| **J12A** | J1 | 22°42'17″N | 109°13'15″E |
| **I41A** | I4 | 22°16'26″N | 109°28'51″E |
| **B28** | B2 | 22°11'23″N | 108°04'57″E |
| **b2** | b2 | 24°19'59″N | 106°11'20″E |
| **cc3** | cc3 | 24°40'57″N | 106°36'52″E |
| **L14** | L1 | 24°34'56″N | 107°04'13″E |
| **X31** | X3 | 24°40'38″N | 109°41'35″E |
| **Y13** | Y1 | 24°41'59″N | 105°16'01″E |
| **Z14** | Z1 | 24°31'10″N | 105°07'16″E |

Table S2. Genome sequencing and mapping statistics of the 48 earthworm samples.

| Sample | Mapped Reads (M) | Mapped Bases (G) | Avg. Depth | Coverage >= 1X (%) | Coverage >= 5X (%) | Mapping Rate (%) |
|---|---|---|---|---|---|---|
| **B28** | 230.06 | 32.98 | 18.79 | 67.91 | 50.60 | 95.26 |
| **b2** | 265.43 | 36.95 | 28.94 | 96.25 | 92.03 | 99.19 |
| **C22A** | 78.72 | 12.04 | 13.75 | 92.64 | 83.42 | 98.18 |
| **C23A** | 75.24 | 12.02 | 12.70 | 93.15 | 83.11 | 94.23 |
| **cc3** | 266.22 | 37.60 | 31.18 | 96.75 | 93.20 | 99.38 |
| **D11A** | 78.11 | 12.02 | 13.59 | 93.05 | 83.68 | 97.60 |
| **D13A** | 79.47 | 12.02 | 13.28 | 92.71 | 83.31 | 99.19 |
| **D22A** | 78.84 | 12.00 | 14.35 | 92.63 | 83.85 | 98.61 |
| **D23A** | 79.30 | 12.03 | 14.17 | 92.55 | 83.73 | 98.95 |
| **E21A** | 79.14 | 12.02 | 13.73 | 92.68 | 82.66 | 98.82 |
| **E22A** | 79.34 | 12.00 | 13.23 | 92.31 | 82.01 | 99.20 |
| **E23A** | 78.22 | 12.03 | 13.28 | 92.34 | 81.91 | 97.70 |
| **E31A** | 77.00 | 11.81 | 13.54 | 92.25 | 82.76 | 97.88 |
| **E41A** | 79.16 | 12.02 | 13.64 | 92.45 | 82.59 | 98.89 |
| **E42A** | 78.87 | 12.04 | 13.83 | 91.98 | 82.24 | 98.34 |
| **E43A** | 79.01 | 12.00 | 13.74 | 92.22 | 82.39 | 98.78 |
| **F21A** | 79.02 | 12.01 | 11.06 | 93.06 | 81.15 | 98.86 |
| **F31A** | 79.28 | 12.00 | 13.64 | 92.78 | 83.20 | 99.14 |
| **G11A** | 77.84 | 12.03 | 13.74 | 92.96 | 83.60 | 97.20 |
| **G12A** | 77.68 | 12.03 | 13.25 | 92.48 | 83.06 | 96.98 |
| **G13A** | 77.22 | 12.02 | 13.59 | 92.33 | 82.92 | 96.54 |
| **G21A** | 75.82 | 12.03 | 12.47 | 93.13 | 83.18 | 94.85 |
| **G22A** | 79.67 | 12.04 | 13.53 | 92.82 | 83.93 | 99.26 |
| **G23A** | 79.13 | 12.02 | 13.29 | 93.44 | 84.23 | 98.79 |
| **H11A** | 79.31 | 12.02 | 13.98 | 92.07 | 83.26 | 99.04 |
| **H12A** | 79.14 | 12.01 | 13.69 | 92.04 | 83.07 | 98.88 |
| **H13A** | 78.62 | 12.05 | 14.00 | 91.83 | 82.93 | 97.99 |
| **H21A** | 78.77 | 12.00 | 13.78 | 92.44 | 83.41 | 98.52 |
| **H22A** | 78.79 | 12.02 | 13.79 | 92.56 | 83.52 | 98.39 |
| **H31A** | 79.17 | 12.01 | 14.07 | 92.38 | 83.32 | 98.91 |
| **H32A** | 79.41 | 12.02 | 14.14 | 92.28 | 83.38 | 99.17 |
| **H33A** | 79.04 | 12.03 | 13.71 | 92.55 | 83.22 | 98.60 |
| **I11A** | 79.13 | 12.03 | 13.28 | 93.33 | 83.45 | 98.78 |
| **I12A** | 74.05 | 12.04 | 12.70 | 93.59 | 83.40 | 93.00 |
| **I13A** | 79.07 | 12.00 | 13.81 | 92.66 | 83.33 | 98.90 |
| **I31A** | 77.98 | 12.02 | 13.29 | 93.24 | 83.01 | 97.44 |
| **I32A** | 79.18 | 12.01 | 14.02 | 93.01 | 83.61 | 98.98 |
| **I33A** | 79.57 | 12.04 | 13.42 | 92.80 | 82.98 | 99.20 |
| **I41A** | 78.39 | 12.02 | 13.62 | 92.57 | 83.22 | 97.95 |
| **I42A** | 78.06 | 12.01 | 13.55 | 92.14 | 82.60 | 97.61 |
| **I43A** | 74.59 | 12.03 | 12.85 | 92.30 | 82.16 | 93.38 |
| **J12A** | 77.35 | 12.03 | 12.89 | 93.50 | 83.57 | 96.81 |
| **J21A** | 77.57 | 12.00 | 12.65 | 92.45 | 82.20 | 97.14 |
| **J22A** | 75.82 | 12.04 | 12.76 | 92.69 | 82.49 | 94.81 |
| **L14** | 183.00 | 32.20 | 14.71 | 53.03 | 38.29 | 84.90 |
| **X31** | 294.25 | 43.20 | 36.24 | 95.43 | 91.32 | 99.33 |
| **Y13** | 259.18 | 39.07 | 34.39 | 81.78 | 71.39 | 98.14 |
| **Z14** | 254.21 | 38.00 | 38.94 | 96.94 | 93.34 | 99.18 |

*Note:*
Mapped reads and bases were calculated using Samtools stats. Coverage was estimated using mosdepth.

Table S3. Summary of genomic variant filtering stages and final datasets for the 48 Asp individuals.

| **Filtering Stage** | **Filtering Criteria** | **Number of SNPs** | **Retention (%)** |
|---|---|---|---|
| Initial Variants | Raw joint calling results across 48 individuals | 112,087,883 | **100%** |
| Hard Filtered & Masked | QUAL > 30, MQ > 40, FS < 60, SOR < 3; Exclude TE regions; Bi-allelic SNPs | 73,599,596 | **65.66%** |
| Final PopGen Set | Genotype DP < 5 set as missing; Missing rate < 10%; MAF > 0.05 | 10,769,051 | **9.61%** |

*Note:*

Variant filtering was executed using BCFtools (v1.22). Repeats were masked based on a TE-specific GFF annotation.

Table S4. Summary of individual heterozygosity rates across the 48 individuals.

| Sample ID | Group | No. of Het Sites | No. of Hom Sites | Total Variants | SNP Het Ratio (%) | Genomic Het Rate (%) |
|---|---|---|---|---|---|---|
| **Group1** | | | | | | |
| E21A | Group1 | 1,877,401 | 5,715,807 | 7,593,208 | 24.7247 | 0.2513 |
| E22A | Group1 | 1,904,299 | 5,736,986 | 7,641,285 | 24.9212 | 0.2549 |
| E23A | Group1 | 1,910,152 | 5,693,755 | 7,603,907 | 25.1207 | 0.2557 |
| E41A | Group1 | 1,884,823 | 5,734,425 | 7,619,248 | 24.7377 | 0.2523 |
| E42A | Group1 | 2,011,196 | 5,872,300 | 7,883,496 | 25.5115 | 0.2693 |
| E43A | Group1 | 1,904,784 | 5,753,549 | 7,658,333 | 24.8720 | 0.2550 |
| F31A | Group1 | 1,779,180 | 5,995,549 | 7,774,729 | 22.8841 | 0.2382 |
| I31A | Group1 | 1,696,567 | 5,575,480 | 7,272,047 | 23.3300 | 0.2271 |
| I32A | Group1 | 1,797,279 | 5,853,638 | 7,650,917 | 23.4910 | 0.2406 |
| I33A | Group1 | 1,771,468 | 5,723,126 | 7,494,594 | 23.6366 | 0.2372 |
| **Group2** | | | | | | |
| C22A | Group2 | 1,718,941 | 6,258,023 | 7,976,964 | 21.5488 | 0.2301 |
| C23A | Group2 | 1,720,372 | 6,137,777 | 7,858,149 | 21.8928 | 0.2303 |
| D11A | Group2 | 1,831,897 | 6,345,328 | 8,177,225 | 22.4024 | 0.2453 |
| D13A | Group2 | 1,868,688 | 6,398,560 | 8,267,248 | 22.6035 | 0.2502 |
| D22A | Group2 | 1,768,651 | 6,276,970 | 8,045,621 | 21.9828 | 0.2368 |
| D23A | Group2 | 1,771,856 | 6,222,544 | 7,994,400 | 22.1637 | 0.2372 |
| G11A | Group2 | 1,667,870 | 6,075,833 | 7,743,703 | 21.5384 | 0.2233 |
| G12A | Group2 | 1,778,104 | 6,220,836 | 7,998,940 | 22.2292 | 0.2380 |
| G13A | Group2 | 1,804,982 | 6,249,712 | 8,054,694 | 22.4091 | 0.2416 |
| G21A | Group2 | 1,653,673 | 6,215,089 | 7,868,762 | 21.0157 | 0.2214 |
| G22A | Group2 | 1,802,242 | 6,417,316 | 8,219,558 | 21.9263 | 0.2413 |
| G23A | Group2 | 1,672,887 | 6,232,160 | 7,905,047 | 21.1623 | 0.2240 |
| X31 | Group2 | 1,948,550 | 6,648,316 | 8,596,866 | 22.6658 | 0.2609 |
| Z14 | Group2 | 1,362,477 | 6,189,702 | 7,552,179 | 18.0408 | 0.1824 |
| b2 | Group2 | 1,511,144 | 5,764,003 | 7,275,147 | 20.7713 | 0.2023 |
| cc3 | Group2 | 1,467,772 | 5,704,990 | 7,172,762 | 20.4631 | 0.1965 |
| **Group3** | | | | | | |
| E31A | Group3 | 2,097,221 | 6,317,511 | 8,414,732 | 24.9232 | 0.2808 |
| F21A | Group3 | 1,952,589 | 6,062,239 | 8,014,828 | 24.3622 | 0.2614 |
| H11A | Group3 | 2,143,556 | 6,536,155 | 8,679,711 | 24.6962 | 0.2870 |
| H12A | Group3 | 2,194,229 | 6,556,345 | 8,750,574 | 25.0753 | 0.2938 |
| H13A | Group3 | 2,173,012 | 6,539,408 | 8,712,420 | 24.9415 | 0.2909 |
| H21A | Group3 | 1,983,447 | 6,379,617 | 8,363,064 | 23.7168 | 0.2655 |
| H22A | Group3 | 2,052,875 | 6,385,008 | 8,437,883 | 24.3293 | 0.2748 |
| H31A | Group3 | 2,031,036 | 6,365,633 | 8,396,669 | 24.1886 | 0.2719 |
| H32A | Group3 | 2,018,679 | 6,383,565 | 8,402,244 | 24.0255 | 0.2703 |
| H33A | Group3 | 1,993,543 | 6,304,769 | 8,298,312 | 24.0235 | 0.2669 |
| I11A | Group3 | 1,823,063 | 5,917,926 | 7,740,989 | 23.5508 | 0.2441 |
| I12A | Group3 | 1,762,229 | 5,748,709 | 7,510,938 | 23.4622 | 0.2359 |
| I13A | Group3 | 1,947,470 | 6,171,914 | 8,119,384 | 23.9854 | 0.2607 |
| I41A | Group3 | 1,900,591 | 6,120,088 | 8,020,679 | 23.6961 | 0.2544 |
| I42A | Group3 | 2,120,535 | 6,282,639 | 8,403,174 | 25.2349 | 0.2839 |
| I43A | Group3 | 1,918,203 | 6,087,063 | 8,005,266 | 23.9618 | 0.2568 |
| J12A | Group3 | 1,845,759 | 5,880,081 | 7,725,840 | 23.8907 | 0.2471 |
| J21A | Group3 | 1,977,692 | 6,305,878 | 8,283,570 | 23.8749 | 0.2648 |
| J22A | Group3 | 1,958,275 | 6,294,655 | 8,252,930 | 23.7282 | 0.2622 |
| **OutGroup** | | | | | | |
| B28 | OutGroup | 2,295,409 | 3,946,467 | 6,241,876 | 36.7743 | 0.3073 |
| L14 | OutGroup | 1,541,970 | 2,328,387 | 3,870,357 | 39.8405 | 0.2064 |

*Note:* "SNP Het Ratio" refers to the percentage of heterozygous sites among identified variants, while "Genomic Het Rate" is calculated based on the estimated genome size of 746.95 Mb.

| Sample ID | Group | No. of Het Sites | No. of Hom Sites | Total Variants | SNP Het Ratio (%) | Genomic Het Rate (%) |
|---|---|---|---|---|---|---|
| Y13 | OutGroup | 2,933,207 | 5,127,729 | 8,060,936 | 36.3879 | 0.3927 |

*Note:* "SNP Het Ratio" refers to the percentage of heterozygous sites among identified variants, while "Genomic Het Rate" is calculated based on the estimated genome size of 746.95 Mb.

Table S5. Summary of population genetic statistics and pairwise Fst.

| Population | n | Pi | Tajima's D | SNP Het Ratio (%) | Fst (vs. G1) | Fst (vs. G2) | Fst (vs. G3) | Fst (vs. OutGroup) |
|---|---|---|---|---|---|---|---|---|
| **Group1** | 10 | 0.0044 | 0.8374 | 24.323 | NA | 0.1045 | 0.1344 | 0.3634 |
| **Group2** | 16 | 0.0042 | 1.0160 | 21.551 | 0.1045 | NA | 0.0835 | 0.3748 |
| **Group3** | 19 | 0.0037 | 0.8297 | 24.193 | 0.1344 | 0.0835 | NA | 0.4324 |

*Note:* G1: Group1; G2: Group2; G3: Group3.

Table S6. Divergence time estimates and 95% HPD intervals based on MCMCtree analysis.

| **Divergence Event** | **Mean Age** | **95% HPD Interval** | **Calibration Type / Prior** |
|---|---|---|---|
| Root: C. teleta vs. Sedentaria/Errantia | 4.703 | [4.084, 5.227] | Fossil constraint B(4.0, 5.2) |
| Split: E. andrei vs. Amynthas clade | 2.301 | [1.994, 2.571] | Fossil constraint B(2.0, 3.6) |
| Speciation: A. aspergillum vs. A. corticis | 0.478 | [0.412, 0.537] | Estimated |

*Note:*
HPD: Highest Posterior Density interval.

Table S7. Inferred effective population size (Ne) history for three earthworm populations using SMC++.

| Population Group | Time (Ma) | Estimated Ne |
|---|---|---|
| **Group1** | 1.6362 | 1.23e+06 |
| **Group1** | 0.3086 | 2.79e+06 |
| **Group1** | 0.1770 | 2.91e+05 |
| **Group2** | 1.3917 | 1.20e+06 |
| **Group2** | 0.0434 | 8.99e+04 |
| **Group3** | 1.4192 | 1.33e+06 |
| **Group3** | 0.0300 | 6.49e+04 |

*Note:*
Time is presented in Million years ago (Ma) from ancient to modern; Ne is presented in scientific notation.
[*] Mutation rate = 1.18e-09; Generation time = 0.6 year.

**Table S8. f-branch (fb) statistics quantifying gene flow between earthworm populations.**

| Tree Branch | Descendants | fb (to Group1) | fb (to Group2) | fb (to Group3) |
|---|---|---|---|---|
| **b4** | **Group2** | 0.0000 | - | - |
| **b5** | **Group3** | 0.3185 | - | - |

*Note:*
The f-branch (fb) statistic quantifies the proportion of the genome derived from introgression.
* '-' indicates that the gene flow path is not testable given the tree topology.

Table S9. Individual summary of Runs of Homozygosity (ROH) and inbreeding coefficients (F_ROH).

| Sample | Group | N_ROH | Total Mb | F_ROH | Short (0.3-1Mb) | Medium (1-5Mb) | Long (>5Mb) |
|---|---|---|---|---|---|---|---|
| **E42A** | **Group1** | 107 | 50.31 | 0.0663 | 105 | 2 | 0 |
| **F31A** | **Group1** | 95 | 38.79 | 0.0511 | 95 | 0 | 0 |
| **E43A** | **Group1** | 64 | 27.55 | 0.0363 | 64 | 0 | 0 |
| **E41A** | **Group1** | 55 | 24.59 | 0.0324 | 54 | 1 | 0 |
| **E21A** | **Group1** | 49 | 20.12 | 0.0265 | 49 | 0 | 0 |
| **E23A** | **Group1** | 44 | 18.35 | 0.0242 | 44 | 0 | 0 |
| **E22A** | **Group1** | 40 | 16.73 | 0.0221 | 40 | 0 | 0 |
| **I32A** | **Group1** | 24 | 8.83 | 0.0116 | 24 | 0 | 0 |
| **I33A** | **Group1** | 23 | 8.24 | 0.0109 | 23 | 0 | 0 |
| **I31A** | **Group1** | 7 | 2.53 | 0.0033 | 7 | 0 | 0 |
| **X31** | **Group2** | 225 | 106.25 | 0.1400 | 222 | 3 | 0 |
| **G13A** | **Group2** | 162 | 80.34 | 0.1059 | 157 | 5 | 0 |
| **D13A** | **Group2** | 165 | 75.04 | 0.0989 | 162 | 3 | 0 |
| **D11A** | **Group2** | 155 | 72.22 | 0.0952 | 153 | 2 | 0 |
| **G22A** | **Group2** | 160 | 71.16 | 0.0938 | 159 | 1 | 0 |
| **C22A** | **Group2** | 115 | 51.82 | 0.0683 | 112 | 3 | 0 |
| **D22A** | **Group2** | 114 | 48.41 | 0.0638 | 112 | 2 | 0 |
| **G12A** | **Group2** | 103 | 45.83 | 0.0604 | 102 | 1 | 0 |
| **C23A** | **Group2** | 104 | 45.69 | 0.0602 | 103 | 1 | 0 |
| **D23A** | **Group2** | 104 | 45.02 | 0.0593 | 101 | 3 | 0 |
| **G21A** | **Group2** | 87 | 35.04 | 0.0462 | 87 | 0 | 0 |
| **G11A** | **Group2** | 66 | 28.13 | 0.0371 | 65 | 1 | 0 |
| **G23A** | **Group2** | 70 | 27.94 | 0.0368 | 70 | 0 | 0 |
| **cc3** | **Group2** | 20 | 7.24 | 0.0095 | 20 | 0 | 0 |
| **Z14** | **Group2** | 13 | 4.96 | 0.0065 | 13 | 0 | 0 |
| **b2** | **Group2** | 11 | 4.38 | 0.0058 | 11 | 0 | 0 |
| **H12A** | **Group3** | 252 | 113.81 | 0.1500 | 247 | 5 | 0 |
| **H11A** | **Group3** | 236 | 100.19 | 0.1320 | 235 | 1 | 0 |
| **E31A** | **Group3** | 214 | 97.65 | 0.1287 | 211 | 3 | 0 |
| **I42A** | **Group3** | 176 | 78.96 | 0.1041 | 172 | 4 | 0 |
| **H13A** | **Group3** | 167 | 70.75 | 0.0932 | 167 | 0 | 0 |
| **H32A** | **Group3** | 160 | 69.61 | 0.0917 | 157 | 3 | 0 |
| **I13A** | **Group3** | 142 | 68.03 | 0.0896 | 137 | 5 | 0 |
| **I43A** | **Group3** | 136 | 60.96 | 0.0803 | 135 | 1 | 0 |
| **H31A** | **Group3** | 140 | 58.76 | 0.0774 | 140 | 0 | 0 |
| **H22A** | **Group3** | 140 | 58.15 | 0.0766 | 140 | 0 | 0 |
| **H21A** | **Group3** | 134 | 55.71 | 0.0734 | 134 | 0 | 0 |
| **H33A** | **Group3** | 130 | 55.62 | 0.0733 | 130 | 0 | 0 |
| **J21A** | **Group3** | 122 | 51.42 | 0.0678 | 120 | 2 | 0 |
| **I41A** | **Group3** | 108 | 44.85 | 0.0591 | 108 | 0 | 0 |
| **F21A** | **Group3** | 85 | 32.35 | 0.0426 | 85 | 0 | 0 |
| **I11A** | **Group3** | 71 | 28.66 | 0.0378 | 71 | 0 | 0 |
| **J22A** | **Group3** | 69 | 26.77 | 0.0353 | 69 | 0 | 0 |
| **I12A** | **Group3** | 47 | 18.30 | 0.0241 | 47 | 0 | 0 |
| **J12A** | **Group3** | 26 | 9.43 | 0.0124 | 26 | 0 | 0 |

*Note:*
F_ROH was calculated based on an estimated genome size of 758.86 Mb.
* Counts represent the number of ROH segments in each length category: Short (0.3–1 Mb), Medium (1–5 Mb), and Long (>5 Mb).

Table S10. Summary of genetic load indicators and functional variant distributions based on outgroup-polarized SNPs.

| Category | Group1 | Group2 | Group3 |
|---|---|---|---|
| **Loss of Function (High Impact)** | | | |
| **LoF (Homozygous)** | 2076.1 ± 104.2 | 2122.1 ± 219.9 | 2468.9 ± 171.5 |
| **LoF (Heterozygous)** | 3740.3 ± 198.5 | 3218.7 ± 377.8 | 2928.2 ± 316.3 |
| **Missense (Moderate Impact)** | | | |
| **Missense (Homozygous)** | 23583.9 ± 1182.9 | 24113.3 ± 2574.3 | 28509.4 ± 2335.2 |
| **Missense (Heterozygous)** | 45778.3 ± 2453.9 | 38939.8 ± 4615.8 | 35859.7 ± 3949.5 |
| **General Statistics** | | | |
| **Total Synonymous** | 70467.1 | 63656.4 | 64306.1 |
| **Mis/Syn Ratio** | 0.9843 | 0.9905 | 1.0010 |

*Note:*

Data represent the counts of derived alleles identified via outgroup polarization (Mean ± SD).

[*] Mis/Syn Ratio indicates the efficiency of purifying selection: (Total Missense) / (Total Synonymous).
